# The impact of distributional assumptions in gene-set and pathway analysis: how far can it go wrong?

**DOI:** 10.1101/2021.02.01.429279

**Authors:** Chi-Hsuan Ho, Yu-Jyun Huang, Ying-Ju Lai, Rajarshi Mukherjee, Chuhsing Kate Hsiao

**Affiliations:** Division of Biostatistics and Data Science, Institute of Epidemiology and Preventive Medicine, National Taiwan University, Taipei, Taiwan; Department of Biostatistics, Harvard University, Boston, USA; Bioinformatics and Biostatistics Core, Center of Genomic Medicine, National Taiwan University, Taipei, Taiwan

**Keywords:** association study, gene expression, gene set analysis, multivariate normality test, pathway analysis

## Abstract

Gene-set analysis (GSA) has been one of the standard procedures for exploring potential biological functions when a group of differentially expressed genes have been derived. The development of its methodology has been an active research topic in recent decades. Many GSA methods, when newly proposed, rely on simulation studies to evaluate their performance with a common implicit assumption that the multivariate expression values are normally distributed. The validity of this assumption has been disputed in several studies but no systematic analysis has been carried out to assess the influence of this distributional assumption. Our goal in this study is not to propose a new GSA method but to first examine if the multi-dimensional gene expression data in gene sets follow a multivariate normal distribution (MVN). Six statistical methods in three categories of MVN tests were considered and applied to a total of twenty-two datasets of expression data from studies involving tumor and normal tissues, with ten signaling pathways chosen as the gene sets. Second, we evaluated the influence of non-normality on the performance of current GSA tools, including parametric and non-parametric methods. Specifically, the scenario of mixture distributions representing the case of different tumor subtypes was considered. Our first finding suggests that the MVN assumption should be carefully dealt with. It does not hold true in many applications tested here. The second investigation of the GSA tools demonstrates that the non-normality does affect the performance of these GSA methods, especially when subtypes exist. We conclude that the use of the inherent multivariate normality assumption should be assessed with care in evaluating new GSA tools, since this MVN assumption cannot be guaranteed and this assumption affects strongly the performance of GSA methods. If a newly proposed GSA method is to be evaluated, we recommend the incorporation of multivariate non-normal distributions or sampling from large databases if available.

## 1. INTRODUCTION

Gene-set analysis (GSA) is one of the standard procedures used in biomedical research when interest lies in the evaluation of certain biological functions, such as pathways, or of the collective effect of a set of genes on disease phenotypes (Hirschhorn 2009; Mooney *et al*. 2014). The development of the methodology for performing GSA has attracted much attention in the biostatistics and bioinformatics communities in recent decades. Findings in GSAs can provide a better understanding of disease mechanisms and identify potential treatment targets, particularly for complex diseases (Subramanian *et al*. 2005; Hirschhorn 2009; Mooney *et al*. 2014). Results from the procedures can save time, cost, and effort in translational studies. Most GSA methods are applied to pathways as a way of evaluating and testing pathway association.

Several review articles have documented the evolution of GSA tools (Ackermann and Strimmer 2009; Maciejewski 2013; de Leeuw *et al*. 2016), with emphasis on their differences, assumptions, and statistical properties; some have also addressed issues that arise in GSA, including the lack of consideration of linkage disequilibrium (or correlation) between genes (Goeman and Buhlmann 2007; Gatti *et al*. 2010; Lin *et al*. 2018), the choice of null hypothesis (competitive or self-contained tests) (Goeman and Buhlmann 2007; Maciejewski 2013; de Leeuw *et al*. 2016), and the influence of the proportion of associated genes in the set (Maciejewski 2013; de Leeuw *et al*. 2016). A few studies have expressed doubts about the distributional assumptions made about the data (Kerr *et al*. 2000; Konishi 2004; Zyla *et al*. 2017). Whether or not transcriptomic data are normally distributed, however, has not been resolved (Kerr *et al*. 2000; Konishi 2004; Zyla *et al*. 2017; de Torrenté *et al*., 2019; Liu *et al*. 2019), despite the fact that it has been noted that heterogeneity in the profiling values may exist due to disease subtypes. No study that we are aware of, however, has investigated the impact of distributional assumptions on the conclusions of GSA, when the gene-set exerts a certain molecular function or activity.

The commonly adopted distributional assumptions for gene expression data are the multivariate normal or log-normal distribution. Despite doubts about this assumption, these distributions are frequently used to generate expression values in simulation studies when evaluating the performance of a newly proposed GSA tool. In other words, many GSA tools inherently assume that the gene expression data are normally or log-normally distributed, or that the GSA tests are robust to the distributional assumptions. Since the normality assumption may not be true or guaranteed (Kerr *et al*. 2000; Konishi 2004; Zyla *et al*. 2017), such simulation-based assessment of GSA tools may be incomprehensive, leading to possibly spurious findings derived in such analysis. This argument applies when the gene-set analyses are used for SNP-set or DNA methylation profiling data (Wang *et al*. 2011; Chang *et al*. 2016; Dong *et al*. 2019), since these data are of discrete and percentage type, respectively.

Whether or not the assumed distribution is appropriate may affect the applicability of GSA methods. Performance, for instance, may be affected if the data deviate from normality. To what extent the change in performance corresponds to the degree of the deviation requires further examination. GSA methods that are mean-based (de Leeuw *et al*. 2016) may be able to guard against this issue if the size of the gene-set, hence the number of genes, is small enough relative to the number of samples for asymptotic normality to apply. For non-parametric GSA tests, it is not yet known how robust they are with respect to distributional assumptions. The goals of this study therefore are to investigate the goodness-of-fit of the multivariate normal distributional for the gene expression data and to evaluate the performance of common GSA methods in response to the deviation of the normality assumption. Our approach to the first goal will be based on the case of multivariate instead of univariate normality tests, and our second goal will incorporate both parametric and non-parametric GSA tools.

### 1.1 Normality test of gene sets

In the first part of this study, we will examine whether the parametric multivariate normal (MVN) distribution is a suitable assumption for gene expression data. To assess if the multi-dimensional data set at hand fits a multivariate normal distribution, many tests have been proposed in the literature and several articles have presented comparisons and reviews of them (Thode 2002; Mecklin and Mundfrom 2004; Mecklin and Mundfrom 2005; Korkmaz *et al*. 2014; Chen and Xia 2019). Among these tests, the Mardia test examines the normality assumption based on skewness and kurtosis (Mardia 1970). It is usually referred to as the Mardia skewness test or Mardia kurtosis test, representing its natural extension of univariate skewness and kurtosis tests to the multivariate case. It is therefore not robust to other types of deviations from normality and its power is unstable against many alternatives (Thode 2002; Mecklin and Mundfrom 2005; Zhou and Shao 2014; Chen and Xia 2019). The Henze-Zirkler (HZ) test evaluates the weighted distance between the observed multi-dimensional data and the multivariate normal distribution (Henze and Zirkler 1990).

The distance is usually defined as the squared difference between two corresponding Fourier transformations (characteristic functions) of the observed and expected observations, and then weighted by a kernel function.

The Royston test (Royston 1992) applies the Shapiro-Wilk (SW) statistic (Shapiro and Wilk 1965), a univariate normality test utilizing variance ratios of the order statistics of observed data, to each coordinate of random vectors, and then combines the marginal statistics. In contrast to this, alternative methods focus on transforming the multivariate data into a univariate value via projection, since it is well known that a *p*-variate random vector **X** is normal if and only if *θ***X** is univariate normal for all *θ* ∈ {*θ* ∈ *R*^*p*^ : ǁθǁ = 1}. Following this rationale, the Fattorini (FA) test (Fattorini 1986; Lee *et al*. 2014) and TN test (Zhou and Shao 2014) were proposed based on the projection of **X** and SW statistics, respectively. Another test not in the previous categories is a nonparametric test, the Energy (EN) test (Székely and Rizzo 2005). It defines the “energy” as a function of the distance between the data and the null multivariate normal distribution, and then examines if this Energy statistic achieves statistical significance.

Based on simulation studies, Mecklin and Mundfrom (2005) recommend the HZ and Royston tests based on their stable type I and II error rates. Korkmaz, Goksuluk, and Zararsiz (2014) suggest the HZ test if the sample size is larger than 100, and the Royston test if smaller than 50. The simulation studies in Székely and Rizzo (2005) compared the Energy test with the HZ and Mardia tests and concluded that the Energy and HZ tests have comparable performance. The TN test performs better than FA and HZ when the alternative hypothesis is a mixture of normal, chi-square or gamma distributions (Zhou and Shao 2014). Note that these results and conclusions are based on simulated observations from lower dimensional distributions and a large ratio of sample size to dimension (Mecklin and Mundfrom 2005; Székely and Rizzo 2005; Korkmaz *et al*. 2014), which may not reflect the case of a gene set containing a large number of genes.

To investigate if multivariate normality fits the gene expression data, we downloaded twenty-two datasets from public websites including the National Center for Biotechnology Information (NCBI) and The Cancer Genome Atlas (TCGA) websites. These gene expression values were from tumor and adjacent normal tissues from cancer patients, or from blood samples from subjects free from cancer. Since gene expression values are influenced by genetic and environmental factors, the data from subjects with similar or the same age, gender, ethnicity, cancer stage and grade were further stratified and selected before analysis. These data sets are then tested to determine if they follow the MVN distribution.

### 1.2 Distributional assumptions and gene-set analysis

The second goal of this study is to compare the performance of several GSA tools with emphasis on the impact of the distributional assumption, based on data generated from multivariate normal and non-normal distributions. The objective of GSA is to examine if the gene-set associates with the phenotype, that is, if the gene-set shows a collective effect on the outcome variable, such as a disease subtype or some continuous phenotypic value. When used for pathway analysis, it is common to categorize GSA methods into three generations: over representation analysis (ORA), functional class scoring (FCS), and pathway-topology (PT) methods.

Several studies have reviewed and compared these different GSA categories (Goeman and Buhlmann 2007; Akermann and Strimmer 2009; Gatti *et al*. 2010; Khatri *et al*. 2012; Maciejewski 2013; de Leeuw *et al*. 2016). The ORA approach is concerned with determining if the differentially expressed (DE) genes are *over represented* in a candidate gene-set/pathway; that is, if the number of DE genes in the set is beyond chance alone.

When this is the goal, an ORA test is carried out based on the hypergeometric distribution, chi-square test, or Fisher’s exact test (Boyle *et al*. 2004). The ORA approach is intuitive and is still a widely used procedure to screen for potential targeted pathways from a list of known ones, thus it is often called a knowledge-based procedure. A major limitation of ORA, however, is that it ignores the relationship among genes in the set of interest and considers them equally important. The FCS approach aims to examine the collective and coordinate effect of the gene-set/pathway. It calculates a score at the pathway level based on a statistic evaluated at each individual component gene. Most FCSs consider the score a weighted sum of the gene-level statistics, where the weight usually represents the interrelationship among the genes. The advantage of FCS is that it allows for the inclusion of genes that may not be significant at the single gene level, the incorporation of the dependence among genes, and the potential to compare contributions of different pathways. More discussion of various FCS methods is in Section 3. The third category, the PT methods, such as SPIA (Draghici *et al* 2007), NetGSA (Shojaie and Michailidis 2009), and NetworkHub (Chang *et al*. 2020), all involve a known topological structure of the genes, and therefore can be applied only if the structure of the set of genes is already determined. This limits the application of PT methods to established biological pathways. Consequently, it becomes difficult to evaluate their performance with simulation studies since the biological structure is required as input in PT methods. Here we consider specifically the GSA tools of the FCS type. In particular, we categorize this group into two different subgroups, parametric and nonparametric GSAs.

The parametric GSA tools were developed mostly based on known statistical distributions and models, including the global test (Goeman *et al*. 2004) in a random-effects model, Hotelling’s statistic (Schafer and Strimmer 2005; Lu *et al*. 2005), and ROAST (Wu *et al*. 2010) in a multivariate normal model, and the pathway activity score (P score) in a logistic regression model (Lin *et al*. 2018). The non-parametric GSA methods mostly use the ranks of gene expression levels to reduce the heterogeneity in the absolute values across different genes. Several popular non-parametric methods are Gene Set Enrichment Analysis (Subramanian *et al*. 2005), the N-statistic (Baringhaus and Franz 2004; Klebanov *et al*. 2007; Glazko and Emmert-Streib 2009), and the Kolmogrove-Smirnov test for the mean vector (KS_mean) and the Kolmogrove-Smirnov test for the covariance (KS_var) matrix (Rahmatallah *et al*. 2017). These GSA tools will be investigated to evaluate the change in their performance in response to different distributional assumptions and various degrees of departure from normality.

## 2. MATERIALS AND METHODS

### 2.1 Multivariate normality tests of gene expression values

#### Data source and data management

Here we consider twenty-two data sets to examine if the normal distribution fits the gene expression data. Among these data sets, eight were downloaded from TCGA and fourteen were from NCBI. Most of them were gene expression data from cancer (breast, colorectal, lung, ovarian, and glioblastoma) patients and only four groups were from smokers (COPD_1 and COPD_2) and non-smokers (COPD_3 and COPD_4). Each data set was processed with standard procedures of quality control and normalization for microarray data. Next, considering that expression levels are affected by age, gender, ethnicity, cancer grade and stage, these downloaded data were stratified according to these factors, if available, and then the groups containing a larger number of samples were selected for the following analyses. For instance, in the first data set (Ni et al. 2010), only breast cancer patients who were in grade II and III and were Malaysian Malays were selected. In the second data set (Sabates-Bellver et al. 2007), the 23 colorectal cancer patients with tumors found in the rectum or sigmoid colon were selected from a study originally containing 32 patients. In the glioblastoma (GBM) study in Cancer Genome Atlas Research Network (2008), only male patients aged beyond 50 and diagnosed with the four subtypes were selected. Table 1 lists the type of cancer or disease, the accession number in the NCBI data portal or TCGA identification, the source of the sample (tissue, cell or peripheral blood), the platform (from Affymetrix GPL96/HG-U133A, GPL570/HG-U133plus2, or Multiple from Affymetrix and Agilent) used for obtaining the expression levels, the sample type (paired or independent), and the number of samples in each group.

**Table 1.** Description of the twenty-two gene expression data sets considered for the multivariate normality tests.

| Case | ID<sup>a</sup> | Source<sup>b</sup> | Platform | Paired<sup>e</sup> | No. |
| --- | --- | --- | --- | --- | --- |
| Breast | GSE15852 | T | GPL96 | Y | 24 |
| Colorectal | GSE8671 | T | GPL570 | Y | 23 |
| Lung_1<sup>f</sup> | GSE7670 | T | GPL96 | Y | 21 |
| Lung_2<sup>f</sup> | GSE19804 | T | GPL570 | Y | 47 |
| Lung_3<sup>f</sup> | GSE19188 | T | GPL570 | N | 25 |
| Lung_4<sup>f</sup> | TCGA-lusc | T | GPL96 | N | 49 |
| Lung_5<sup>f</sup> | TCGA-lusc | T | GPL96 | N | 44 |
| Ovarian_1<sup>c</sup> | TCGA-ov | M | GPL96 | N | 47 |
| Ovarian_2<sup>c</sup> | TCGA-ov | M | GPL96 | N | 116 |
| GBM_c<sup>g</sup> | TCGA-gbm | T | Multiple | N | 66 |
| GBM_m<sup>g</sup> | TCGA-gbm | T | Multiple | N | 78 |
| GBM_n<sup>g</sup> | TCGA-gbm | T | Multiple | N | 47 |
| GBM_p<sup>g</sup> | TCGA-gbm | T | Multiple | N | 58 |
| **Control** |  |  |  |  |  |
| Breast | GSE15852 | T | GPL96 | Y | 24 |
| Colorectal | GSE8671 | T | GPL570 | Y | 23 |
| Lung_1<sup>f</sup> | GSE7670 | T | GPL96 | Y | 21 |
| Lung_2<sup>f</sup> | GSE19804 | T | GPL570 | Y | 47 |
| Lung_3<sup>f</sup> | GSE19188 | T | GPL570 | N | 41 |
| COPD_1<sup>d</sup> | GSE11906 | T | GPL570 | N | 31 |
| COPD_2<sup>d</sup> | GSE42057 | B | GPL570 | N | 27 |
| COPD_3<sup>d</sup> | GSE11906 | T | GPL570 | N | 26 |
| COPD_4<sup>d</sup> | GSE42057 | B | GPL570 | N | 22 |

<sup>a</sup>: GEO accession number or TCGA identification; <sup>b</sup>: T for tissue or cell; B for peripheral blood; M for tissue or blood; <sup>c</sup>: the two groups are poor (Ovarian\_1) versus good (Ovarian\_2) prognosis group; <sup>d</sup>: the two groups (COPD\_1, COPD\_2) are smokers and the two groups (COPD\_3, COPD\_4) are nonsmokers; <sup>e</sup>: if the data are paired samples, then the case group contains cancer tissue and control contains the adjacent normal tissue; <sup>f</sup>: Lung\_1 is non-small cell lung carcinoma (NSCLC) patients with adenocarcinoma, Lung\_2 and Lung\_3 only specify NSCLC, Lung\_4 and Lung\_5 are NSCLC patients with squamous cell carcinoma; <sup>g</sup>: the four groups are the different subtypes, GBM\_c for classical, GBM\_m for mesenchymal, GBM\_n for neural, and GBM\_p for proneural, respectively.

**Table 2.** The number of the rejection rates *Q* larger than 10%, #{*Q* ≥10%}, among the 220 tests (each MVN was tested on 10 pathways for all 22 datasets), and the correlation matrix of the rejection rates *Q* among the six MVN tests.

|  | Energy | HZ | Royston | FA | TN | Mardia |
| --- | --- | --- | --- | --- | --- | --- |
| $\#\{Q \geq 10\}$ | 209 | 213 | 219 | 207 | 220 | 217 |
| % | 99.5% | 96.8% | 99.5% | 94.1% | 100% | 98.6% |
| Correlation matrix |  |  |  |  |  |  |
| Energy | 1 | 0.91 | 0.97 | 0.94 | 0.99 | 0.97 |
| HZ |  | 1 | 0.85 | 0.90 | 0.89 | 0.90 |
| Royston |  |  | 1 | 0.89 | 0.96 | 0.92 |
| FA |  |  |  | 1 | 0.96 | 0.98 |
| TN |  |  |  |  | 1 | 0.99 |
| Mardia |  |  |  |  |  | 1 |

#### Selected statistical methods and gene-sets for MVN tests

Six methods in three categories are considered for the multivariate normality test. The first category includes two tests based on the distance measure between the *p*-dimensional data and the *p*-variate normal distribution, one is the EN and the other is the HZ test. The distance measure used in EN is the Euclidean distance and in HZ it is the expected distance between two characteristic functions. The second category contains three tests which are all multivariate extensions of the univariate SW test: the Royston test which combines the SW statistic from each of the *p* coordinates, and the FA and TN tests which are based on projection of the multivariate data into a one-dimensional point, where FA and TN utilize different statistics of the projected values. The test in the third group is the traditional Mardia test based on multivariate skewness or kurtosis. We consider any violation in either one as a deviation from multivariate normality and therefore the Mardia test was applied here to test if either the skewness or kurtosis test reached significance at the 5% nominal level.

All the tests were carried out in R, with the function *mvnorm.etest* in the R package *energy* for the one (multivariate) sample energy EN test (Székely and Rizzo 2017), with the function *mvn* in the R package *MVN* for the HZ, Royston, and Mardia tests (Korkmaz *et al*. 2014) and with the functions *faTest* and *mvnTest* in the R package *mvnormalTest* for the Fattorini FA and TN tests (Lee *et al*. 2014), respectively.

#### Selected gene-sets for MVN tests

For the gene sets to be tested for multivariate normality, ten signaling pathways from the Kyoto Encyclopedia of Genes and Genomes (KEGG) were selected deliberately. These signaling pathways were constructed based on interacting molecules involving specific biological functions, where the gene nodes in the pathway network (gene-set) would be expected to exert some correlation. Such pathways targeted by gene-set analysis therefore are the focus of our examination of the normality test. The ten signaling pathways defined in KEGG are the p53, mTOR, Jak-STAT, PI3k-Akt, Wnt, ErbB, MAPK, RAS, TGF-β, and TNF pathways, which contain 60, 69, 32, 85, 76, 47, 115, 69, 58, and 82 gene nodes, respectively. These have been reported to associate with one or more of the cancers considered here. For example, the p53 pathway was reported to show association with breast (Gasco *et al.* 2002; Bertheau *et al*. 2008; Walerych *et al.* 2012), colorectal (Li *et al.* 2015), lung (Mitsudomi *et al.* 2000; Shtivelman *et al.* 2014), ovarian (Bernardini *et al.* 2010; Hayano *et al.* 2014), and glioblastoma (Cancer Genome Atlas Research Network 2008; Jung *et al.* 2019) cancers. Association of the other pathways such as mTOR, Jak-STAT, PI3k-Akt, Wnt, RAS, and TGF-β have been reported as well (Park and Kim 2007; Cancer Genome Atlas Research Network 2008; Eroles *et al.* 2012; Shtivelman *et al.* 2014; Slattery *et al.* 2014; Lin *et al*. 2018; Jung *et al*. 2019). For COPD studies, the pathways reported to associate with COPD or smoking status include Jak-STAT (Bahr *et al*. 2013; Yew-Booth *et al*. 2015; Nicholson *et al*. 2016), PI3k-Akt (Marwick *et al*. 2010; Kim *et al*. 2011; Bahr *et al*. 2013; Mercado *et al*. 2015), and mTor (Bahr *et al*. 2013; Mercado *et al*. 2015).

### 2.2 Evaluating the impact of the distributional assumption

#### Parametric and non-parametric GSA tools

The first group of GSAs contains four parametric tests: the global test (Goeman et al. 2004), Hotelling’s statistic (Schafer and Strimmer 2005; Lu et al. 2005), ROAST (Wu et al. 2010), and the pathway activity score (P score) in the logistic regression model (Lin et al. 2018). The global test examines the existence of heterogeneity among the random gene effects under a parametric generalized linear model. The function gt in the R package Globaltest performs this test. The Hotelling’s T^2^ statistic compares the mean vectors of the expression profiles from subjects of different disease statuses with a modified estimate of the covariance matrix to incorporate possible correlation among genes. This test can be performed in the R package Hotelling with the function hotelling.test. The ROAST method is also constructed under the linear model. It combines the gene-level modified t-statistic to formulate a statistic for the gene-set, and determines the p-value, not via permutation of genes or samples, but by the rotation of the independent residual space to incorporate the intergenic correlations (Langsrud 2005). The function roast in the R package limma performs this test. The pathway activity score, P score, is a statistic based on ranks of gene expression values and magnitudes of correlations (Lin et al. 2018). The score is computed for each sample and the enrichment analysis for association with the phenotype is then tested in a logistic regression model under the case-control study design.

The second group contains non-parametric GSA tests which utilize mostly the ranks of the expression values or some metric of the distance between genes. The first one we considered is the popular gene set enrichment analysis (GSEA) modified for the self-contained null hypothesis for a predetermined set of genes (Gentleman *et al*. 2008). It is nonparametric in the sense that no distributional assumption is made for the expression value and the test statistic is based on the ranks of the degree of association between the phenotype and the individual gene. It assumes independence among gene expression profiles and utilizes rankings and non-parametric distance metrics to derive p values. This test can be carried out with the function *gseattperm* in the R package *Category*. The second test, usually called the N-statistic, compares the Euclidean distance between multivariate observations from two phenotypic groups (Baringhaus and Franz 2004; Klebanov *et al*. 2007; Glazko and Emmert-Streib 2009). It follows the same rationale as in the energy test in previous sections for MVN tests, where a group of observations is compared against observations generated from a MVN distribution. Here for GSA, the focus is on the comparison between two groups of multivariate observations without distributional assumptions for either group. This test can be conducted with the function *eqdist.etest* in the R package *energy* (Székely and Rizzo 2017). The other two GSA tools considered here first rank the multivariate samples based on the minimum spanning tree and then use the multivariate Kolmogorov-Smirnov (KS) test under the self-contained null hypothesis to test for the difference in mean (KS_mean), or in variance (Rahmatallah *et al*. 2017). The R package adopted here is *GSAR* and the functions are *KStest* and *RKStest*, respectively. All the software and functions used in this study are listed in Table 3.

**Table 3.**
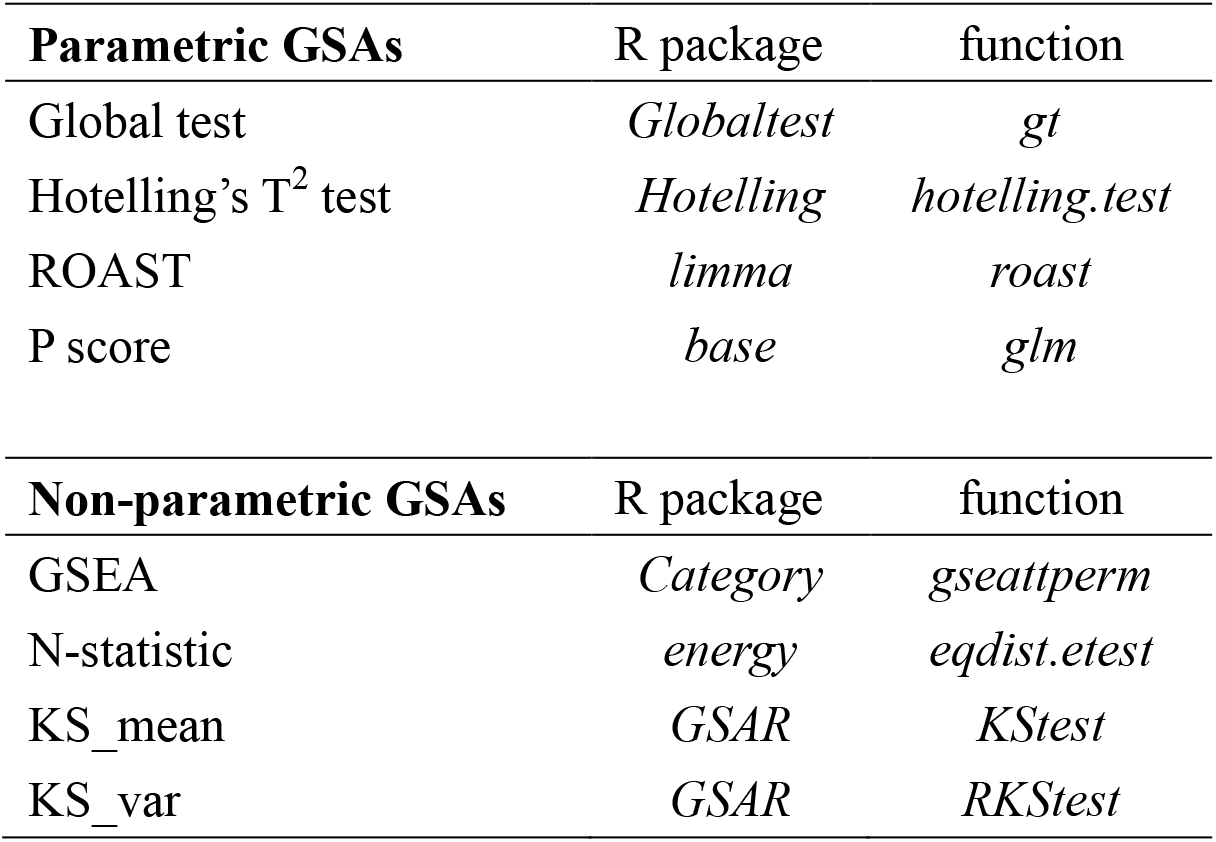
Software (R package; function) used for assessing the performance of the GSA tools.

| **Parametric GSAs** | R package | function |
| --- | --- | --- |
| Global test | *Globaltest* | *gt* |
| Hotelling's $T^2$ test | *Hotelling* | *hotelling.test* |
| ROAST | *limma* | *roast* |
| P score | *base* | *glm* |
| **Non-parametric GSAs** | R package | function |
| GSEA | *Category* | *gseattperm* |
| N-statistic | *energy* | *eqdist.etest* |
| KS_mean | *GSAR* | *KStest* |
| KS_var | *GSAR* | *RKStest* |

#### Simulation setting I, Single component distribution per group (settings A and B)

To evaluate the performance of these GSA tools with respect to different distributions of the gene expression values, the following scenarios were considered. The first two are single component models. Simulation setting A was designed under the *p*-dimensional multivariate normality assumption (*p*=30), where both phenotypic groups, termed as *case* and *control* with 50 subjects in each group, were generated from MVN distributions with different mean vectors (the difference is denoted as 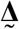) and the same compound symmetry (CS) covariance matrix (with *ρ* = 0.1).

Phenotype group 1 (*control*) and 2 (*case*) in setting A:

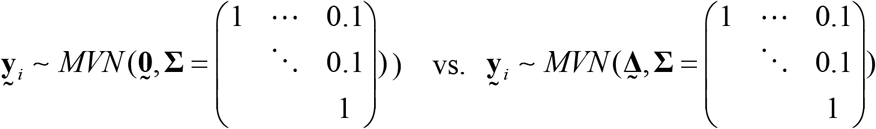

Setting B aims for a larger heterogeneity than A and the data were instead generated from multivariate t distributions (MVT) with 3 degrees of freedom, the same mean vectors as in A and CS covariance matrix with *ρ* = 0.5.

#### Simulation setting II, Mixture of two distributions per group (settings C and D)

In this setting, we consider distributions containing more variation than the above scenario by assuming a mixture of two distributions. In setting C, the expression values were randomly generated from a mixture of two MVN distributions, each with 50% weight. Specifically, in the *control* group, the mean vectors of the two component MVNs are the zero mean vector 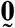 and the vector of all-ones 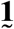, respectively; while in the *case* group, the two mean vectors are 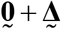 and 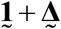, respectively.

Phenotype group 1 (*control*) in setting C:

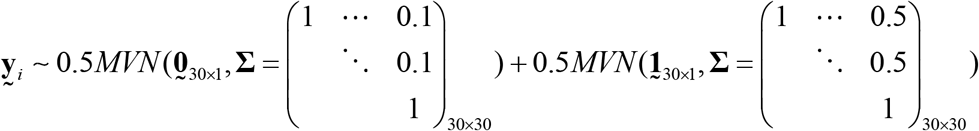

In setting D, the variability in each component is made even larger by assuming a MVT distribution with 3 degrees of freedom.

#### Simulation setting III, When case group contains subtypes (settings E, F, G, & H)

In contrast to the previous scenarios where both *case* and *control* groups are from the same distribution but with different means, here we consider the scenario where the disease (case) group contains more than one component distribution. These situations can arise due to disease subtypes such as the cancer grade, staging, and tumor tissue types (Kim *et al*. 2010). Examples include lung cancer with two major subtypes, small cell lung cancer (SCLC) and non-small cell lung cancer (NSCLC), and leukemia with four major subtypes, acute lymphoblastic leukemia (ALL), acute myeloid leukemia (AML), chronic myeloid leukemia (CML), and chronic lymphocytic leukemia (CLL). The therapeutic choice may differ according to different subtypes where such classification is usually performed clinically with immunohistochemistry (IHC). Therefore, in settings E and F, the distribution for the *case* group is a mixture of two normal (setting E) and two t (setting F) distributions, respectively; whereas in settings G and H, it is a 3-component mixture distribution. Table 4 lists all the distributions considered in these simulation studies.

**Table 4.**
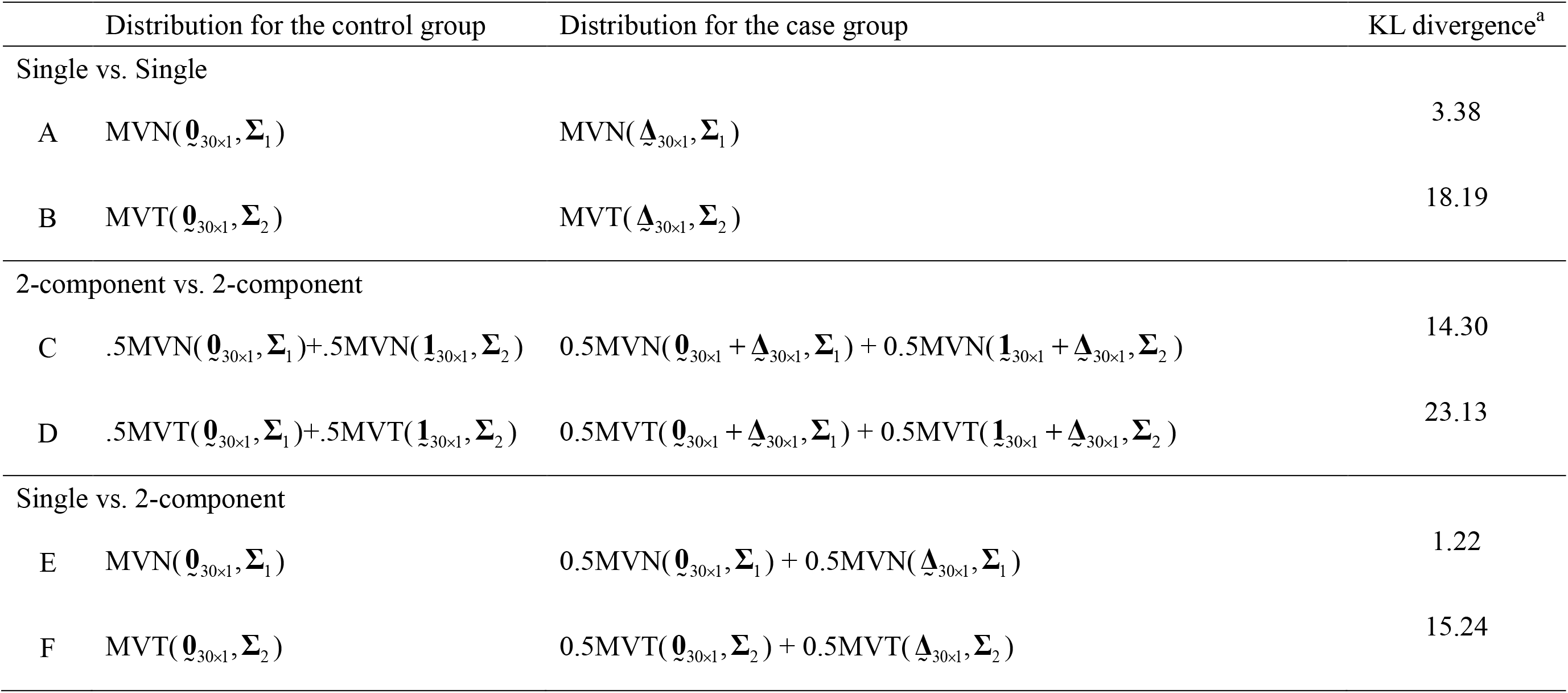
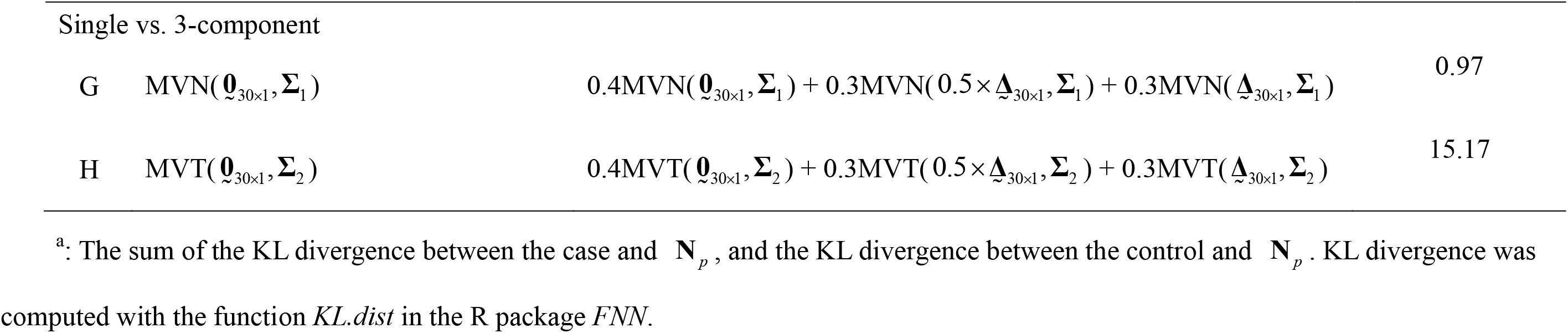
Distributions used to generate expression data in the GSA analysis. MVN stands for the multivariate normal distribution and MVT stands for multivariate t distribution. The two covariance matrices adopted both have compound symmetry correlation structure where the correlation is 0.1 in **Σ**_1_ and 0.5 in **Σ**_2_, respectively. The Kullback-Leibler (KL) divergence measures the distance between the distribution based on simulated data and the MVN with zero mean vector and an identity covariance matrix.

The use of mixture distributions for tumor subtypes incorporates the scenario of skewed distributions which are apparently non-normally distributed (Kim *et al.* 2010). To demonstrate the degree of deviation from normality, the KL divergence is adopted to measure the *distance* between the distribution of the simulated data and **N**_*p*_, a MVN distribution with a zero mean vector and an identity covariance matrix, in each setting. The values are listed in the right-most column in Table 4. Larger values of KL imply a greater difference between the group with the simulated distribution and the group with a MVN distribution.

### 2.3 Data and Software Availability

All the human gene expression data included in this study can be freely downloaded from the NCBI or TCGA website. GEO accession number or TCGA identification were listed in Table 1. The R code for the four multivariate normality tests can be downloaded from https://github.com/r05849032/Four_MVN_tests.

## 3. RESULTS

### 3.1 Normality tests of gene expression values

Since multivariate normality will not hold true for the distribution of any pathway if any subset of the pathway deviates from the normality assumption, we selected a subset of either 10 (if the number of tissue samples exceeds 30) or 5 (if the number is below 30) genes from each pathway and carried out the above six MVN tests. This procedure was repeated 1000 times and the proportion 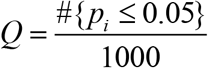 of replications in which the p-value was less than the nominal 0.05 was recorded. These proportions are presented in tables in the online supplementary materials (Tables S1–S6).

For each of the MVN tests, the majority of the *Q*’s (across data sets and pathways) are much greater than 10% as shown in the boxplots in Figure 1 and the top row of Table 2, indicating noticeable evidence against the normality assumption. Next, the average of the *Q*’s across the ten signaling pathways for each data set was calculated, since the magnitudes of the proportions are similar across pathways. These averages are presented in the heatmap in Figure 2. The figure again concludes that the similarity in rejecting the MVN null hypothesis is apparent among the six tests. The corresponding average values are in the supplementary materials (Table S7). In addition, these proportions are similar across the six tests, as indicated by the large correlations (0.85~0.90) in Table 2 and the correlation plot in Figure 3a. When comparing the tests against the energy (EN) test, the scatter plot in Figure 3b reveals that the Royston test tends to reject more often (93%) than EN; while both the HZ and Mardia tests are more conservative than EN (99% and 98%, respectively). As for the FA and TN tests, their performance is relatively closer to that of the EN test.

**Figure 1.**
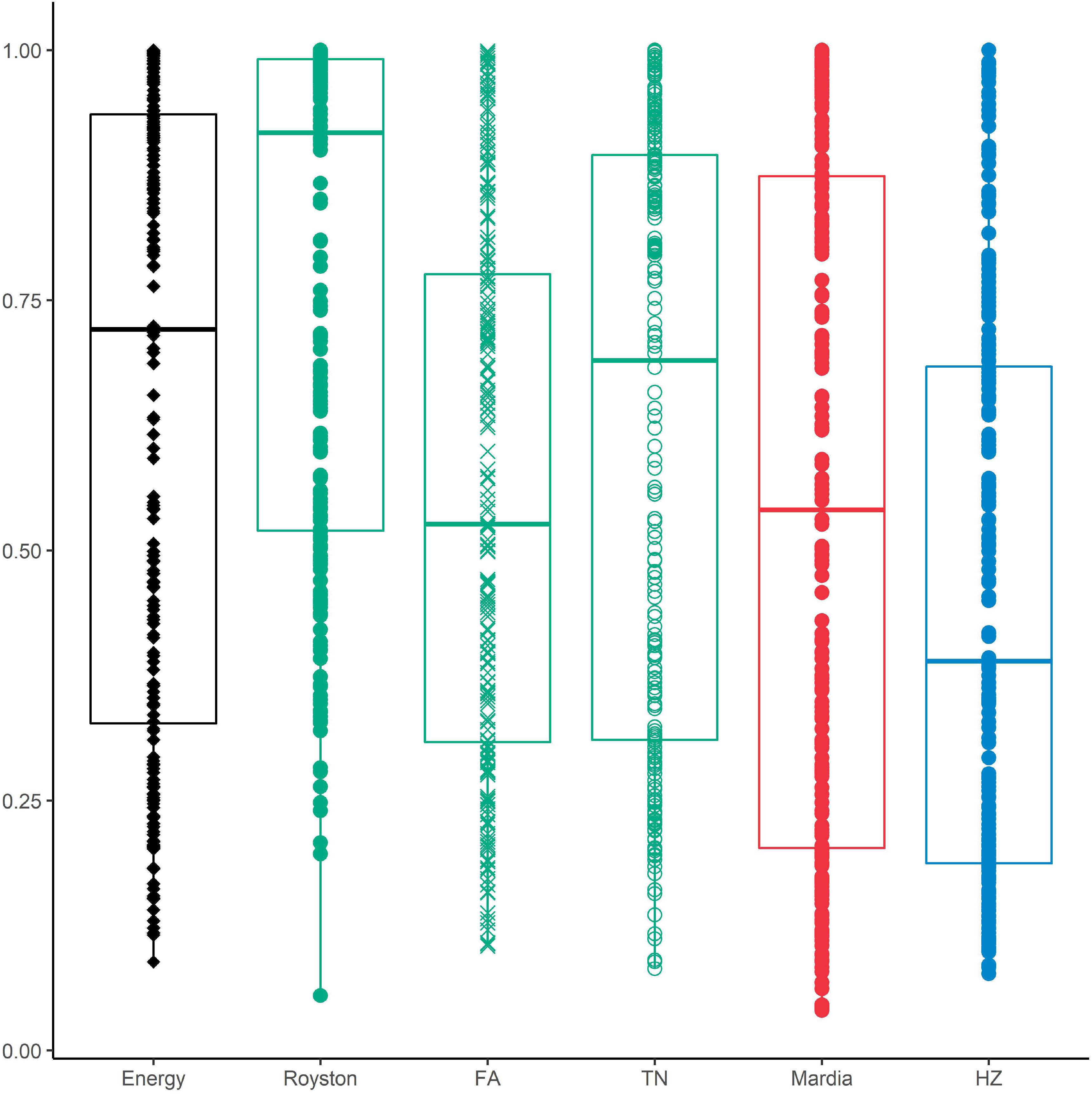
Results of the MVN tests. Boxplots of the 220 proportions of p-values less than 5% under each MVN test; 220 is the product of 22 (datasets) and 10 (pathways).

**Figure 2.**
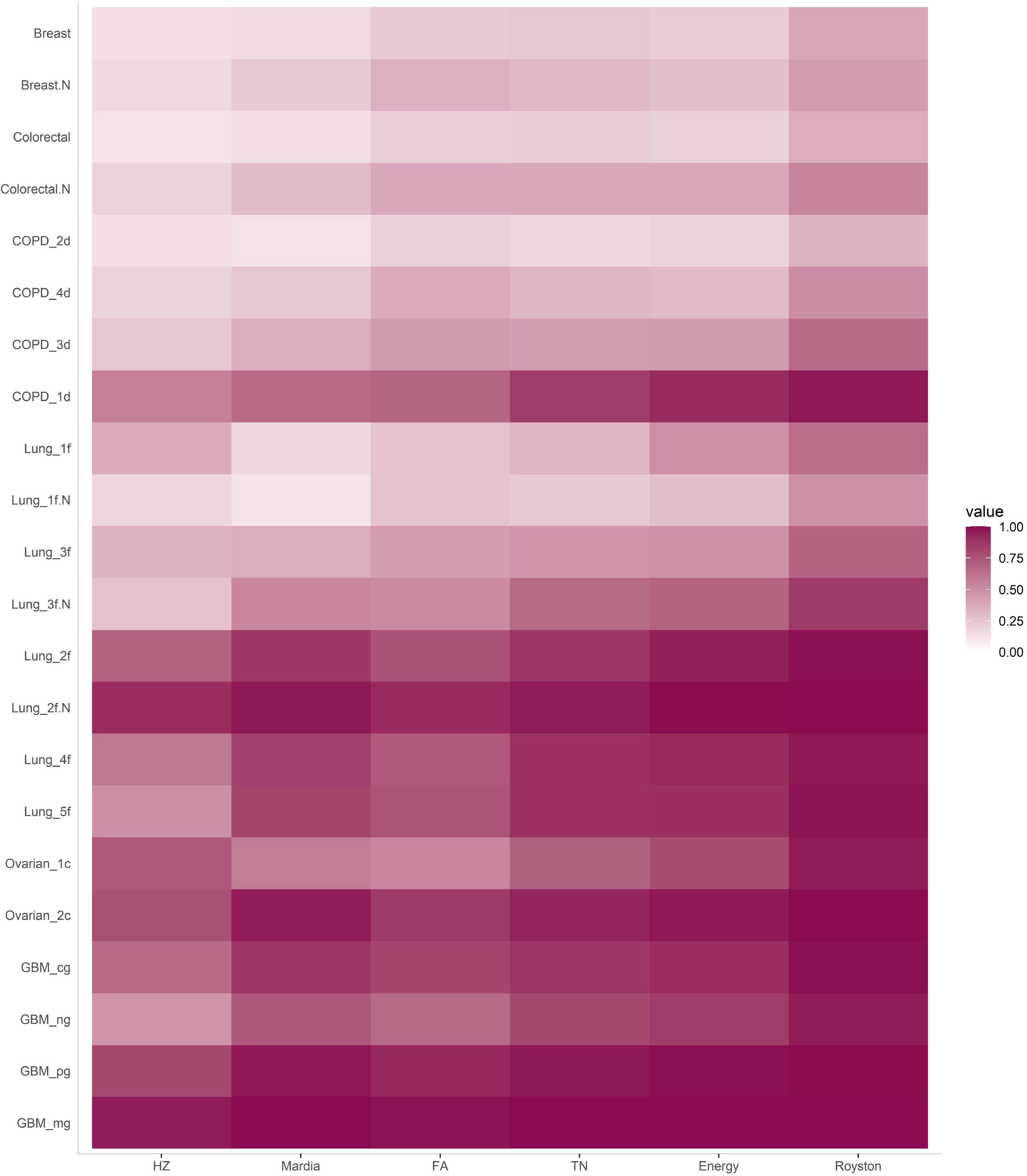
Heatmap of the average proportions across pathways. There are 22 values in each column in the heatmap.

**Figure 3.**
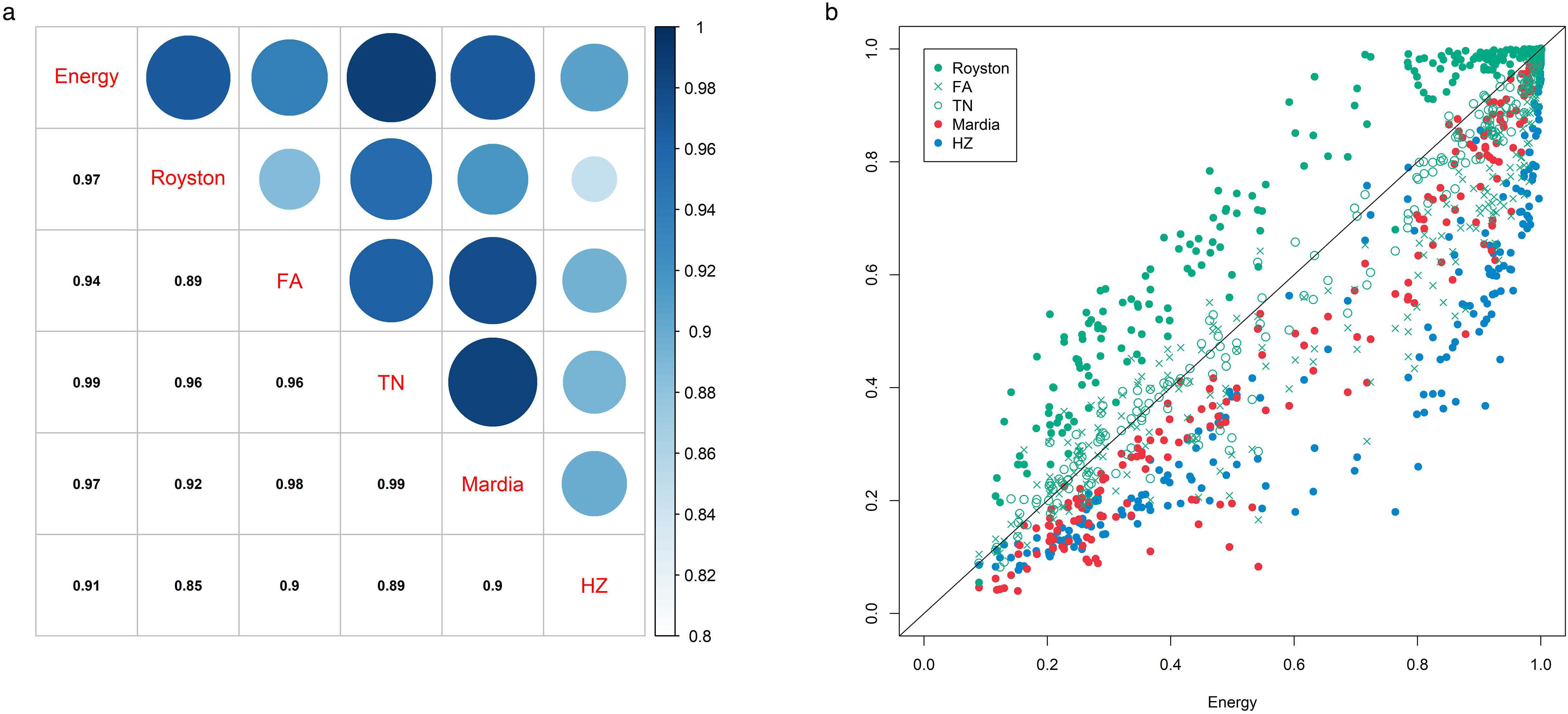
**a** The correlation among these proportions. **b** The scatter plot of these proportions.

### 3.2 Evaluating the impact of the distributional assumption

It has been observed in previous sections that most gene-sets/pathways are not normally distributed, at least not in the applications we have tested. The next task would then be to examine to what degree this distributional assumption affects the performance of current gene set analysis tools.

In each of the three simulation settings, the difference in the mean is denoted as 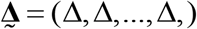, where Δ is set at 0 to evaluate the false positive rate and set at 0.1, 0.3, 0.5, 0.7, and 0.9 to evaluate its power performance. A total of 1000 replications were carried out under each setting and each replication contained 50 *control* subjects and 50 disease (*case*) subjects. The results are shown in Figure 4 and the supplementary materials (Tables S8–S15).

**Figure 4.**
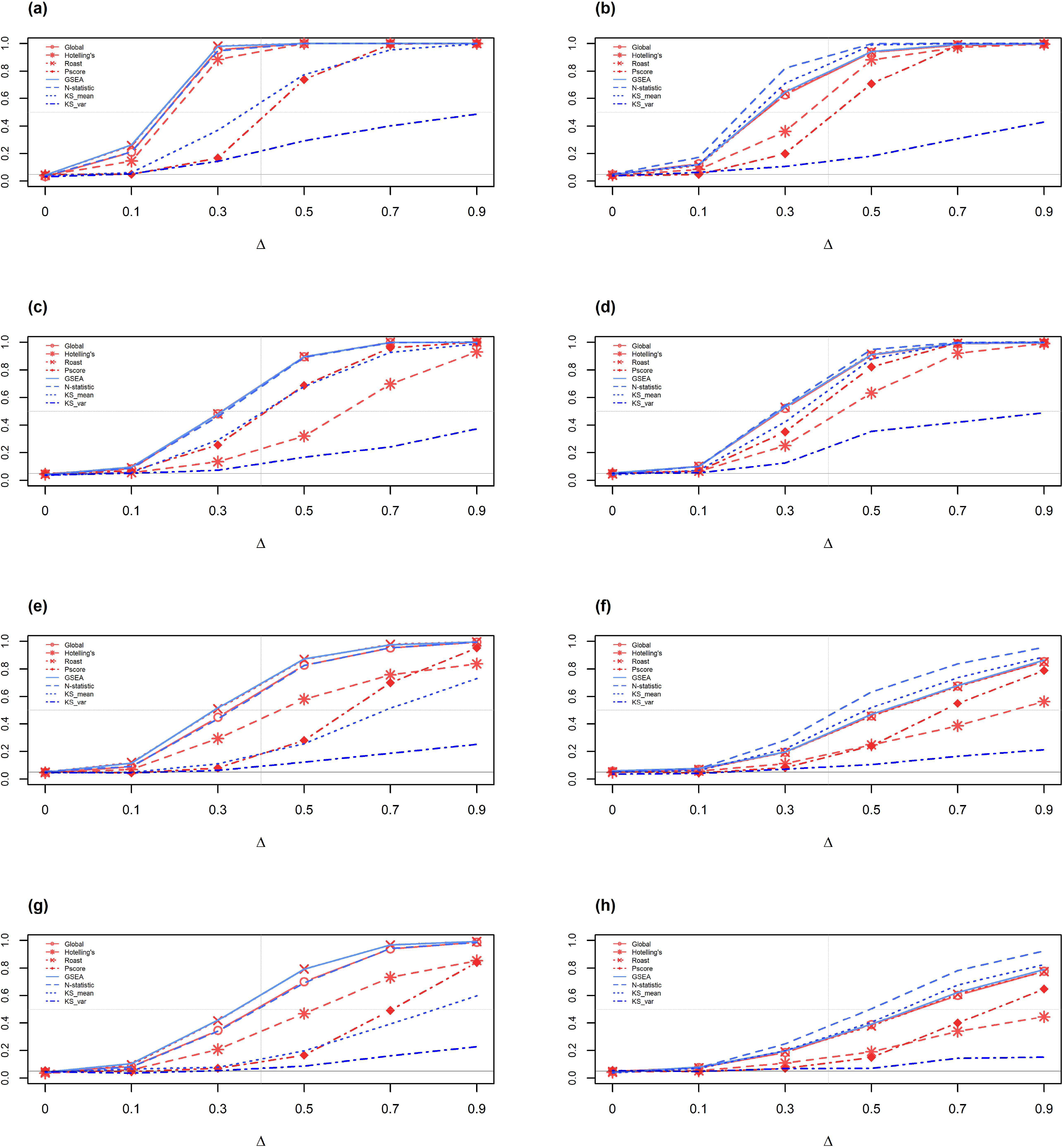
Performance evaluation of eight GSA tests. **a** Setting A, single MVN vs. single MVN. **b** Setting B, single MVT vs. single MVT. **c** Setting C, 2-component mixture MVN vs. 2-component mixture MVN. **d** Setting D, 2-component mixture MVT vs. component mixture MVT. **e** Setting E, single MVN vs. 2-component mixture MVN. **F** Setting F, single MVT vs. 2-component mixture MVT. **g** Setting G, single MVN vs. component mixture MVN. **h** Setting H, single MVT vs. 3-component mixture MVT.

Generally, all tests maintain a reasonable type I error rate at the nominal level 0.05, and their power elevates as the difference Δ increases. However, when normality is no longer valid, the power curve starts to subside. First, note that when comparing A versus B, C vs. D, E vs. F, and G vs. H in Figure 4, the power performance under the MVT, single or mixture components, is always worse than that with MVN as the component distribution. The Kullback-Leibler (KL) divergence in Table 4 also implies that, as compared with the settings (A, C, E, G) in the left panel in Figure 4, the settings (B, D, F, H) in the right panel present larger deviation from MVN. Second, when looking at the left panel in Figure 4, though they all use MVN as the components, the distributions are becoming more skewed due the increasing number of mixture components. Therefore, the power lines become less steep. Third, the drop in power is more obvious in the right panel, when the component distribution is the MVT. In other words, when the disease subtypes are considered as a mixture of distributions, these GSA tools will not perform as well as when the gene expression follows a single component normal distribution.

Among these GSA tools, the global test and ROAST in the category of parametric GSAs and the GSEA and N-statistic in the non-parametric category perform better than the others. These four tools steadily outperform the rest, with the N-statistic standing out when the component distributions are MVT rather than MVN. In other words, the N-statistic is less sensitive to the normality assumptions than the other GSA tools. If we focus only on the MVT (right panel in Figure 4), the performance of KS_mean is as good as or better than GSEA.

## 4. DISCUSSION

The goals of this study are first to investigate whether the multivariate normal distribution is a suitable assumption for gene expression data, and second to examine the performance of common GSA methods in response to deviations from normality. When testing the normality assumption on twenty-two real gene expression data sets, six tests in three categories were considered, including two tests (EN and HZ) based on distance measures, three tests (Royston, FA, and TN) that are multivariate extensions of the univariate Shapiro-Wilk test, and the Mardia test of either multivariate skewness or kurtosis. All these tests indicated strong evidence for rejecting the MVN normality assumption. In a recent research study (de Torrenté *et al*. 2019), based on the shape of gene expression data, many distributions other than the normal were fitted, though in a one-dimensional manner. Similar findings of the non-MVN normality of gene expression data have been concluded in Ho’s research (2018). In addition, we computed the KL divergence between the breast cancer data (GSE15852) in Table 1 and **N**_*p*_ (here *p* varies according to the dimension of the pathway), for each of the ten pathways considered in Section 2. The values are all much larger than zero, indicating a substantial departure from MVN (Table 5).

**Table 5.** The KL divergence between the pathway gene expression (breast cancer data, GSE15852) and the MVN **N**_*p*_, where *p* stands for the number of genes in the corresponding pathway.

| Pathway | No. of genes | Normal vs. $\mathbf{N}_p$ | Tumor vs. $\mathbf{N}_p$ |
| --- | --- | --- | --- |
| ERBB | 45 | 259.57 | 256.91 |
| Jak-Stat | 30 | 164.36 | 165.99 |
| MAPK | 108 | 614.04 | 606.42 |
| mTOR | 57 | 322.89 | 316.87 |
| P53 | 53 | 296.34 | 287.63 |
| PI3K-AKT | 75 | 419.63 | 412.77 |
| RAS | 64 | 373.07 | 369.85 |
| TGF-BETA | 51 | 285.61 | 277.81 |
| TNF | 71 | 401.24 | 394.22 |
| Wnt | 66 | 374.10 | 362.50 |

Based on this non-normality observation, we then proceeded to examine the influence of deviation from normality on the performance of several popular GSA methods. The simulation studies show that, even with a single component MVT distribution, the larger variability can cause a loss in power. When the number of components increases, corresponding to a greater degree of skewness and possibly multimodality of the distribution, the loss becomes worse. We should bear in mind that the GSA tools may not be as good as we wish them to be.

Among the GSA tools examined here, a non-parametric GSA, the N-statistic, steadily performed better than the others. Therefore, this test is recommended when one is uncertain about the distributional assumption; otherwise, applying more than one GSA tool should reach the same conclusion for association study. In addition, when a new GSA tool is proposed and simulation studies are designed to demonstrate its performance, we advocate the incorporation of non-normal distributions, such as the mixture distributions considered here. Although the MVN assumption has been questioned in previous research, to our knowledge, this is the first comprehensive investigation focusing on this issue, as well as the first to compare the performance of GSAs in non-normal scenarios.

There are other issues in the development of GSAs requiring attention. First, since current statistical models may not describe properly the gene expression data under study (de Torrenté *et al*. 2019), a remedy for the simulation design would be to simulate from available databases where the sample size is big enough to cover the heterogeneity across individuals to serve as a pseudo-population. This solution may not be readily put into practice now, but it is not entirely infeasible. As long as data accumulation continues at a high speed and access remains open to users, such a choice should be available in the near future. This argument applies similarly to other genomic data such as RNA-seq and SNP data. Second, the GSA tools we consider here are limited to the FCS category. Currently, many new GSA methods are developed based on given pathway topology (Draghici *et al*. 2007; Shojaie and Michailidis 2009; Chang *et al*. 2020). Most of these tools do not incorporate probabilistic randomness and statistical models, thus it is not easy to investigate their performance using statistical simulation studies to test their robustness against distributional assumptions. Effort would be required by the statistical community to formulate procedures for such evaluation. Finally, current GSA methods treat the gene-set or pathway as a deterministic group of genes. No uncertainty is considered, nor quantified.

However, when the content of a pathway is retrieved from different pathway databases, it is quite possible that the derived pathways are different. Future GSA should incorporate this difference, either by taking the intersection or union of these component nodes, or by including a mechanism such as latent variables for this difference. More studies are clearly warranted.

### Article Summary

The validity of the multivariate normal distribution (MVN) for gene expression data analysis has been examined in this study, and the results indicate that it is very unlikely that the multi-dimensional data follows such distribution. It is further demonstrated that some GSA tools suffer statistical power loss if data do not follow MVN. When proposing a new GSA method, both normal and non-normal distributed expression data should be considered for applications, to guard against the possible failure in reproducibility.

## Declarations

## Ethics approval and consent to participate

Not applicable. All analyses were performed either on publicly available data or on simulation data

## Consent for publication

Not applicable.

## Authors’ contributions

CHH, CKH conceived the study and designed the method. RM designed the normality tests. CHH, YJH, YJL implemented the data analysis and simulation studies. HCC and CKH prepared the draft. All authors read and approved the final manuscript.

## Acknowledgments

The authors thank Dr. Yongzhao Shao and Dr. Ming Zhou of the Division of Biostatistics, New York University School of Medicine for sharing the R code to carry out the TN test. This work was supported in part by the Taiwan Ministry of Science and Technology (MOST-106-2314-B-002-097-MY3; MOST 109-2314-B-002-152).

## Supplementary information

Supplementary materials are available and include Tables for MVN tests (Table S1–S6), average proportions across ten pathways for each dataset and test (Table S7), and for GSA evaluation (Table S8–S15). The R code for the four multivariate normality tests can be downloaded from https://github.com/r05849032/Four_MVN_tests.

## Figure legends

**Table S1.** Energy Test: These are the proportions *Q*, 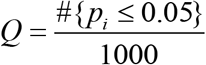, of p values less than the nominal 0.05 among 1000 replications with respect to each data set (row) and pathway (column).

|  | P53 | mTOR | Jak-STAT | PI3k-Akt | Wnt | ERBB | MAPK | RAS | TGF_BETA | TNF |
| --- | --- | --- | --- | --- | --- | --- | --- | --- | --- | --- |
| Breast | 0.216 | 0.162 | 0.247 | 0.155 | 0.322 | 0.256 | 0.219 | 0.209 | 0.118 | 0.290 |
| Colorectal | 0.203 | 0.205 | 0.204 | 0.204 | 0.224 | 0.227 | 0.153 | 0.243 | 0.250 | 0.167 |
| Lung_1f | 0.687 | 0.367 | 0.282 | 0.480 | 0.532 | 0.542 | 0.499 | 0.495 | 0.445 | 0.441 |
| Lung_2f | 0.981 | 0.934 | 0.940 | 0.960 | 0.978 | 0.986 | 0.928 | 0.966 | 0.939 | 0.861 |
| Lung_3f | 0.359 | 0.427 | 0.489 | 0.464 | 0.507 | 0.477 | 0.413 | 0.469 | 0.655 | 0.480 |
| Lung_4f | 0.929 | 0.923 | 0.915 | 0.969 | 0.917 | 0.978 | 0.837 | 0.799 | 0.950 | 0.922 |
| Lung_5f | 0.825 | 0.913 | 0.912 | 0.891 | 0.900 | 0.910 | 0.810 | 0.934 | 0.935 | 0.908 |
| Ovarian_1c | 0.795 | 0.718 | 0.715 | 0.785 | 0.895 | 0.870 | 0.784 | 0.592 | 0.902 | 0.724 |
| Ovarian_2c | 0.924 | 0.951 | 0.982 | 0.969 | 0.981 | 0.995 | 0.981 | 0.983 | 0.994 | 0.987 |
| GBM_cg | 0.974 | 0.865 | 0.851 | 0.860 | 0.867 | 0.944 | 0.885 | 0.873 | 0.917 | 0.973 |
| GBM_mg | 1.000 | 1.000 | 1.000 | 1.000 | 0.999 | 1.000 | 0.999 | 0.999 | 1.000 | 0.997 |
| GBM_ng | 0.910 | 0.842 | 0.955 | 0.825 | 0.817 | 0.862 | 0.803 | 0.811 | 0.847 | 0.785 |
| GBM_pg | 0.999 | 0.992 | 1.000 | 0.987 | 0.987 | 0.971 | 0.981 | 0.997 | 0.995 | 0.978 |
| Breast.N | 0.233 | 0.320 | 0.141 | 0.347 | 0.336 | 0.345 | 0.290 | 0.206 | 0.398 | 0.234 |
| Colorectal.N | 0.311 | 0.367 | 0.541 | 0.346 | 0.329 | 0.463 | 0.353 | 0.450 | 0.235 | 0.365 |
| Lung_1f.N | 0.286 | 0.264 | 0.271 | 0.263 | 0.267 | 0.434 | 0.278 | 0.336 | 0.267 | 0.152 |
| Lung_2f.N | 0.992 | 0.989 | 0.990 | 0.997 | 0.998 | 0.999 | 0.996 | 0.998 | 0.997 | 0.973 |
| Lung_3f.N | 0.633 | 0.602 | 0.932 | 0.698 | 0.548 | 0.764 | 0.631 | 0.801 | 0.702 | 0.554 |
| COPD_1d | 0.857 | 0.952 | 0.920 | 0.921 | 0.878 | 0.839 | 0.908 | 0.922 | 0.968 | 0.926 |
| COPD_2d | 0.227 | 0.257 | 0.089 | 0.116 | 0.182 | 0.130 | 0.210 | 0.123 | 0.267 | 0.253 |
| COPD_3d | 0.353 | 0.507 | 0.395 | 0.431 | 0.389 | 0.416 | 0.490 | 0.294 | 0.468 | 0.616 |
| COPD_4d | 0.256 | 0.395 | 0.183 | 0.251 | 0.287 | 0.202 | 0.381 | 0.545 | 0.282 | 0.286 |

**Table S2.** HZ Test: These are the proportions *Q*, 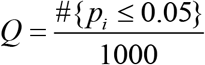, of p values less than the nominal 0.05 among 1000 replications with respect to each data set (row) and pathway (column).

|  | P53 | mTOR | Jak-STAT | PI3k-Akt | Wnt | ERBB | MAPK | RAS | TGF_BETA | TNF |
| --- | --- | --- | --- | --- | --- | --- | --- | --- | --- | --- |
| Breast | 0.144 | 0.084 | 0.135 | 0.085 | 0.175 | 0.158 | 0.130 | 0.118 | 0.112 | 0.172 |
| Colorectal | 0.104 | 0.129 | 0.103 | 0.101 | 0.152 | 0.130 | 0.077 | 0.118 | 0.128 | 0.107 |
| Lung_1f | 0.554 | 0.272 | 0.206 | 0.350 | 0.417 | 0.505 | 0.384 | 0.393 | 0.323 | 0.293 |
| Lung_2f | 0.787 | 0.640 | 0.616 | 0.753 | 0.744 | 0.768 | 0.598 | 0.711 | 0.738 | 0.481 |
| Lung_3f | 0.211 | 0.308 | 0.347 | 0.329 | 0.387 | 0.338 | 0.264 | 0.268 | 0.468 | 0.349 |
| Lung_4f | 0.654 | 0.638 | 0.611 | 0.712 | 0.601 | 0.705 | 0.450 | 0.353 | 0.610 | 0.599 |
| Lung_5f | 0.389 | 0.549 | 0.521 | 0.531 | 0.499 | 0.368 | 0.388 | 0.450 | 0.571 | 0.557 |
| Ovarian_1c | 0.678 | 0.758 | 0.661 | 0.790 | 0.838 | 0.696 | 0.734 | 0.563 | 0.856 | 0.706 |
| Ovarian_2c | 0.650 | 0.639 | 0.817 | 0.670 | 0.817 | 0.887 | 0.721 | 0.689 | 0.895 | 0.859 |
| GBM_cg | 0.847 | 0.512 | 0.667 | 0.635 | 0.605 | 0.762 | 0.545 | 0.548 | 0.652 | 0.774 |
| GBM_mg | 0.988 | 0.981 | 1.000 | 0.986 | 0.954 | 0.979 | 0.940 | 0.958 | 0.974 | 0.875 |
| GBM_ng | 0.674 | 0.363 | 0.572 | 0.489 | 0.507 | 0.375 | 0.454 | 0.356 | 0.454 | 0.418 |
| GBM_pg | 0.981 | 0.792 | 0.946 | 0.775 | 0.795 | 0.690 | 0.700 | 0.735 | 0.854 | 0.682 |
| Breast.N | 0.129 | 0.183 | 0.099 | 0.199 | 0.204 | 0.188 | 0.140 | 0.107 | 0.232 | 0.134 |
| Colorectal.N | 0.171 | 0.183 | 0.274 | 0.159 | 0.196 | 0.200 | 0.188 | 0.222 | 0.147 | 0.191 |
| Lung_1f.N | 0.157 | 0.157 | 0.154 | 0.184 | 0.192 | 0.237 | 0.200 | 0.172 | 0.160 | 0.123 |
| Lung_2f.N | 0.846 | 0.904 | 0.901 | 0.934 | 0.946 | 0.968 | 0.924 | 0.977 | 0.897 | 0.781 |
| Lung_3f.N | 0.293 | 0.180 | 0.611 | 0.253 | 0.186 | 0.180 | 0.216 | 0.260 | 0.277 | 0.226 |
| COPD_1d | 0.471 | 0.609 | 0.530 | 0.605 | 0.545 | 0.390 | 0.514 | 0.565 | 0.748 | 0.567 |
| COPD_2d | 0.122 | 0.168 | 0.086 | 0.083 | 0.111 | 0.122 | 0.135 | 0.099 | 0.212 | 0.197 |
| COPD_3d | 0.210 | 0.264 | 0.191 | 0.243 | 0.236 | 0.195 | 0.245 | 0.141 | 0.313 | 0.414 |
| COPD_4d | 0.114 | 0.245 | 0.134 | 0.161 | 0.172 | 0.109 | 0.254 | 0.382 | 0.200 | 0.174 |

**Table S3.** Royston Test: These are the proportions *Q*, 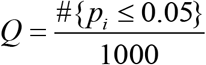, of p values less than the nominal 0.05 among 1000 replications with respect to each data set (row) and pathway (column).

|  | P53 | mTOR | Jak-STAT | PI3k-Akt | Wnt | ERBB | MAPK | RAS | TGF_BETA | TNF |
| --- | --- | --- | --- | --- | --- | --- | --- | --- | --- | --- |
| Breast | 0.338 | 0.264 | 0.451 | 0.279 | 0.551 | 0.405 | 0.347 | 0.401 | 0.240 | 0.535 |
| Colorectal | 0.366 | 0.264 | 0.530 | 0.334 | 0.320 | 0.490 | 0.264 | 0.340 | 0.449 | 0.248 |
| Lung_1f | 0.809 | 0.574 | 0.511 | 0.649 | 0.740 | 0.614 | 0.600 | 0.560 | 0.664 | 0.642 |
| Lung_2f | 0.998 | 0.987 | 0.990 | 0.998 | 0.993 | 0.997 | 0.977 | 0.984 | 0.980 | 0.975 |
| Lung_3f | 0.598 | 0.611 | 0.713 | 0.639 | 0.709 | 0.749 | 0.646 | 0.701 | 0.810 | 0.685 |
| Lung_4f | 0.953 | 0.984 | 0.963 | 0.992 | 0.984 | 0.996 | 0.965 | 0.951 | 0.973 | 0.991 |
| Lung_5f | 0.975 | 0.981 | 0.993 | 0.982 | 0.984 | 0.985 | 0.923 | 0.985 | 0.995 | 0.963 |
| Ovarian_1c | 0.961 | 0.867 | 0.990 | 0.969 | 0.987 | 0.981 | 0.976 | 0.906 | 0.994 | 0.986 |
| Ovarian_2c | 0.998 | 0.999 | 1.000 | 0.996 | 1.000 | 1.000 | 0.998 | 0.999 | 1.000 | 0.999 |
| GBM_cg | 0.995 | 0.992 | 0.996 | 0.988 | 0.983 | 0.998 | 0.971 | 0.968 | 0.990 | 1.000 |
| GBM_mg | 1.000 | 1.000 | 1.000 | 1.000 | 1.000 | 1.000 | 0.999 | 0.999 | 1.000 | 0.998 |
| GBM_ng | 0.984 | 0.981 | 0.99 | 0.911 | 0.912 | 0.952 | 0.982 | 0.984 | 0.924 | 0.991 |
| GBM_pg | 1.000 | 0.996 | 1.000 | 0.996 | 0.982 | 0.993 | 0.991 | 0.993 | 0.991 | 0.998 |
| Breast.N | 0.330 | 0.470 | 0.392 | 0.543 | 0.502 | 0.459 | 0.504 | 0.320 | 0.519 | 0.283 |
| Colorectal.N | 0.374 | 0.610 | 0.715 | 0.548 | 0.446 | 0.784 | 0.448 | 0.617 | 0.351 | 0.548 |
| Lung_1f.N | 0.572 | 0.437 | 0.520 | 0.533 | 0.364 | 0.603 | 0.409 | 0.557 | 0.514 | 0.264 |
| Lung_2f.N | 0.998 | 0.995 | 0.997 | 1.000 | 0.999 | 1.000 | 0.997 | 0.997 | 1.000 | 0.986 |
| Lung_3f.N | 0.951 | 0.851 | 0.984 | 0.900 | 0.713 | 0.680 | 0.847 | 0.936 | 0.930 | 0.760 |
| COPD_1d | 0.941 | 0.957 | 0.994 | 0.976 | 0.997 | 0.988 | 0.968 | 0.963 | 0.991 | 0.972 |
| COPD_2d | 0.481 | 0.486 | 0.055 | 0.208 | 0.328 | 0.340 | 0.400 | 0.197 | 0.421 | 0.435 |
| COPD_3d | 0.495 | 0.744 | 0.541 | 0.654 | 0.666 | 0.672 | 0.717 | 0.575 | 0.658 | 0.793 |
| COPD_4d | 0.494 | 0.491 | 0.455 | 0.443 | 0.487 | 0.355 | 0.521 | 0.678 | 0.505 | 0.456 |

**Table S4.**
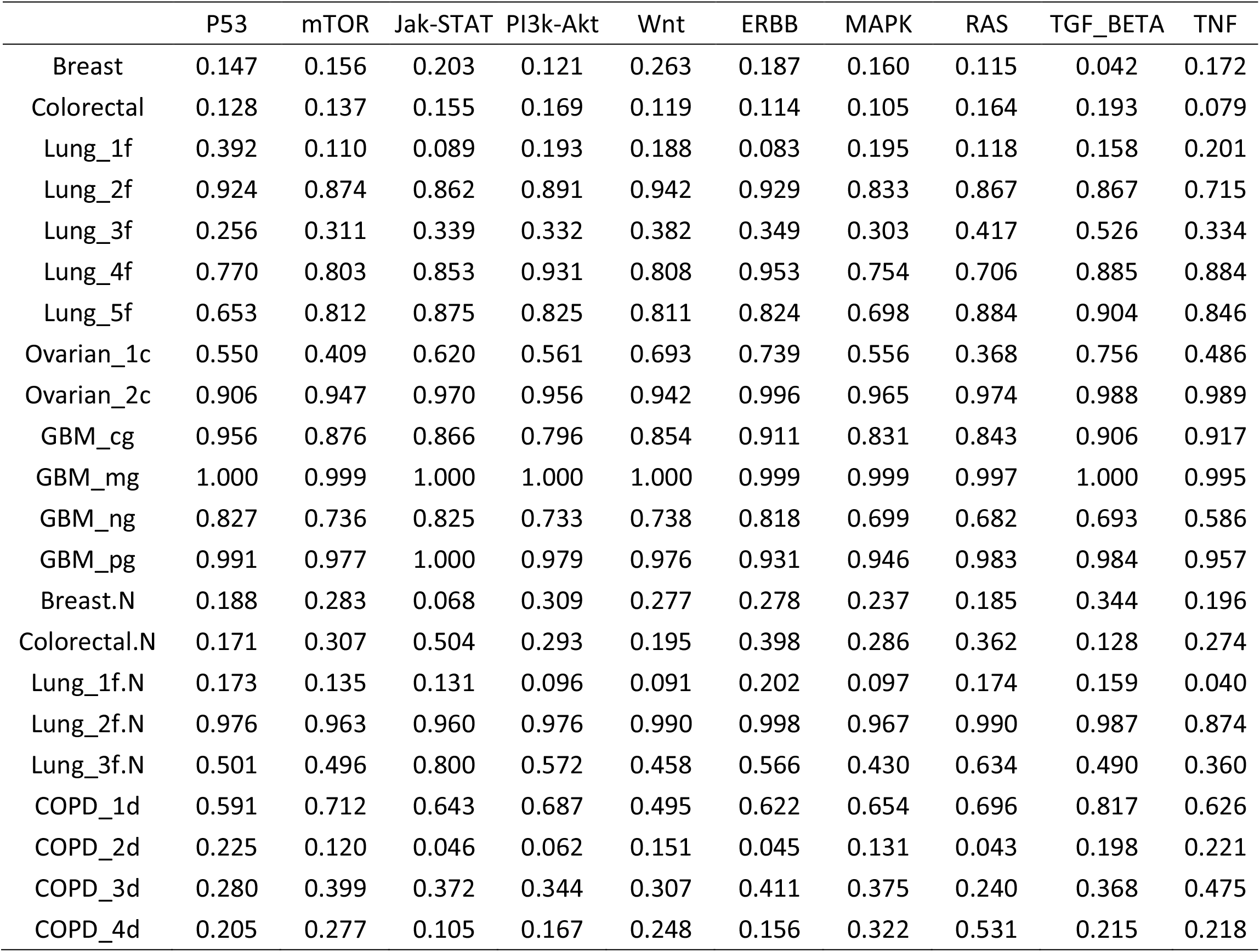
Mardia Test: These are the proportions *Q*, 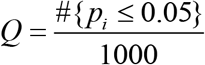, of p values less than the nominal 0.05 among 1000 replications with respect to each data set (row) and pathway (column).

**Table S5.** FA Test: These are the proportions *Q*, 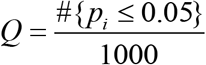, of p values less than the nominal 0.05 among 1000 replications with respect to each data set (row) and pathway (column).

|  | P53 | mTOR | Jak-STAT | PI3k-Akt | Wnt | ERBB | MAPK | RAS | TGF_BETA | TNF |
| --- | --- | --- | --- | --- | --- | --- | --- | --- | --- | --- |
| Breast | 0.236 | 0.290 | 0.280 | 0.159 | 0.387 | 0.279 | 0.212 | 0.181 | 0.108 | 0.278 |
| Colorectal | 0.212 | 0.217 | 0.226 | 0.242 | 0.189 | 0.189 | 0.172 | 0.250 | 0.297 | 0.121 |
| Lung_1f | 0.506 | 0.201 | 0.280 | 0.252 | 0.291 | 0.166 | 0.281 | 0.209 | 0.206 | 0.310 |
| Lung_2f | 0.793 | 0.729 | 0.722 | 0.813 | 0.832 | 0.867 | 0.710 | 0.790 | 0.764 | 0.527 |
| Lung_3f | 0.297 | 0.347 | 0.399 | 0.387 | 0.466 | 0.504 | 0.398 | 0.511 | 0.572 | 0.368 |
| Lung_4f | 0.647 | 0.713 | 0.725 | 0.772 | 0.789 | 0.906 | 0.623 | 0.543 | 0.790 | 0.774 |
| Lung_5f | 0.581 | 0.747 | 0.858 | 0.753 | 0.718 | 0.753 | 0.635 | 0.834 | 0.852 | 0.741 |
| Ovarian_1c | 0.434 | 0.305 | 0.653 | 0.499 | 0.682 | 0.683 | 0.525 | 0.398 | 0.709 | 0.410 |
| Ovarian_2c | 0.738 | 0.774 | 0.924 | 0.808 | 0.868 | 0.964 | 0.765 | 0.886 | 0.973 | 0.891 |
| GBM_cg | 0.888 | 0.855 | 0.867 | 0.718 | 0.782 | 0.888 | 0.685 | 0.769 | 0.866 | 0.794 |
| GBM_mg | 0.992 | 0.987 | 0.999 | 0.995 | 0.974 | 0.989 | 0.967 | 0.964 | 0.997 | 0.955 |
| GBM_ng | 0.719 | 0.551 | 0.735 | 0.660 | 0.711 | 0.623 | 0.656 | 0.671 | 0.725 | 0.455 |
| GBM_pg | 0.979 | 0.913 | 0.986 | 0.961 | 0.918 | 0.859 | 0.834 | 0.928 | 0.957 | 0.887 |
| Breast.N | 0.303 | 0.350 | 0.158 | 0.410 | 0.452 | 0.330 | 0.360 | 0.311 | 0.454 | 0.276 |
| Colorectal.N | 0.217 | 0.399 | 0.526 | 0.405 | 0.246 | 0.551 | 0.420 | 0.467 | 0.181 | 0.321 |
| Lung_1f.N | 0.321 | 0.249 | 0.369 | 0.229 | 0.204 | 0.305 | 0.236 | 0.333 | 0.254 | 0.138 |
| Lung_2f.N | 0.918 | 0.909 | 0.899 | 0.942 | 0.964 | 0.942 | 0.898 | 0.939 | 0.952 | 0.741 |
| Lung_3f.N | 0.469 | 0.449 | 0.626 | 0.574 | 0.471 | 0.505 | 0.446 | 0.599 | 0.468 | 0.438 |
| COPD_1d | 0.641 | 0.704 | 0.696 | 0.727 | 0.539 | 0.669 | 0.657 | 0.707 | 0.807 | 0.656 |
| COPD_2d | 0.358 | 0.191 | 0.105 | 0.132 | 0.183 | 0.128 | 0.166 | 0.107 | 0.298 | 0.288 |
| COPD_3d | 0.357 | 0.523 | 0.468 | 0.411 | 0.397 | 0.471 | 0.389 | 0.296 | 0.500 | 0.560 |
| COPD_4d | 0.321 | 0.422 | 0.228 | 0.288 | 0.378 | 0.304 | 0.443 | 0.642 | 0.329 | 0.354 |

**Table S6.** TN Test: These are the proportions *Q*, 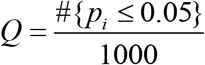, of p values less than the nominal 0.05 among 1000 replications with respect to each data set (row) and pathway (column).

|  | P53 | mTOR | Jak-STAT | PI3k-Akt | Wnt | ERBB | MAPK | RAS | TGF_BETA | TNF |
| --- | --- | --- | --- | --- | --- | --- | --- | --- | --- | --- |
| Breast | 0.195 | 0.202 | 0.317 | 0.177 | 0.393 | 0.277 | 0.238 | 0.220 | 0.117 | 0.320 |
| Colorectal | 0.226 | 0.177 | 0.303 | 0.230 | 0.231 | 0.228 | 0.161 | 0.261 | 0.270 | 0.157 |
| Lung_1f | 0.532 | 0.239 | 0.251 | 0.324 | 0.379 | 0.287 | 0.311 | 0.289 | 0.298 | 0.311 |
| Lung_2f | 0.922 | 0.889 | 0.875 | 0.902 | 0.937 | 0.938 | 0.843 | 0.883 | 0.837 | 0.752 |
| Lung_3f | 0.359 | 0.410 | 0.453 | 0.459 | 0.479 | 0.491 | 0.410 | 0.529 | 0.590 | 0.473 |
| Lung_4f | 0.841 | 0.894 | 0.892 | 0.947 | 0.865 | 0.981 | 0.801 | 0.772 | 0.932 | 0.891 |
| Lung_5f | 0.799 | 0.874 | 0.913 | 0.903 | 0.884 | 0.915 | 0.800 | 0.933 | 0.947 | 0.918 |
| Ovarian_1c | 0.715 | 0.582 | 0.742 | 0.697 | 0.804 | 0.812 | 0.683 | 0.502 | 0.805 | 0.604 |
| Ovarian_2c | 0.848 | 0.908 | 0.975 | 0.925 | 0.931 | 0.974 | 0.941 | 0.947 | 0.961 | 0.945 |
| GBM_cg | 0.940 | 0.849 | 0.881 | 0.802 | 0.843 | 0.896 | 0.850 | 0.832 | 0.873 | 0.929 |
| GBM_mg | 1.000 | 1.000 | 1.000 | 1.000 | 0.999 | 0.998 | 0.993 | 0.995 | 1.000 | 0.985 |
| GBM_ng | 0.864 | 0.803 | 0.895 | 0.782 | 0.727 | 0.844 | 0.799 | 0.779 | 0.742 | 0.708 |
| GBM_pg | 0.991 | 0.974 | 1.000 | 0.980 | 0.976 | 0.923 | 0.942 | 0.985 | 0.976 | 0.945 |
| Breast.N | 0.236 | 0.324 | 0.203 | 0.373 | 0.345 | 0.358 | 0.368 | 0.226 | 0.411 | 0.237 |
| Colorectal.N | 0.248 | 0.397 | 0.622 | 0.395 | 0.282 | 0.519 | 0.347 | 0.478 | 0.221 | 0.372 |
| Lung_1f.N | 0.313 | 0.245 | 0.230 | 0.212 | 0.191 | 0.311 | 0.200 | 0.293 | 0.227 | 0.136 |
| Lung_2f.N | 0.959 | 0.934 | 0.963 | 0.978 | 0.993 | 0.990 | 0.956 | 0.984 | 0.990 | 0.887 |
| Lung_3f.N | 0.634 | 0.658 | 0.858 | 0.718 | 0.559 | 0.642 | 0.556 | 0.769 | 0.704 | 0.490 |
| COPD_1d | 0.795 | 0.853 | 0.853 | 0.855 | 0.798 | 0.807 | 0.851 | 0.847 | 0.950 | 0.853 |
| COPD_2d | 0.255 | 0.211 | 0.089 | 0.112 | 0.196 | 0.091 | 0.184 | 0.082 | 0.250 | 0.264 |
| COPD_3d | 0.342 | 0.466 | 0.405 | 0.434 | 0.408 | 0.416 | 0.438 | 0.315 | 0.424 | 0.563 |
| COPD_4d | 0.295 | 0.363 | 0.202 | 0.285 | 0.355 | 0.200 | 0.383 | 0.513 | 0.304 | 0.309 |

**Table S7:**
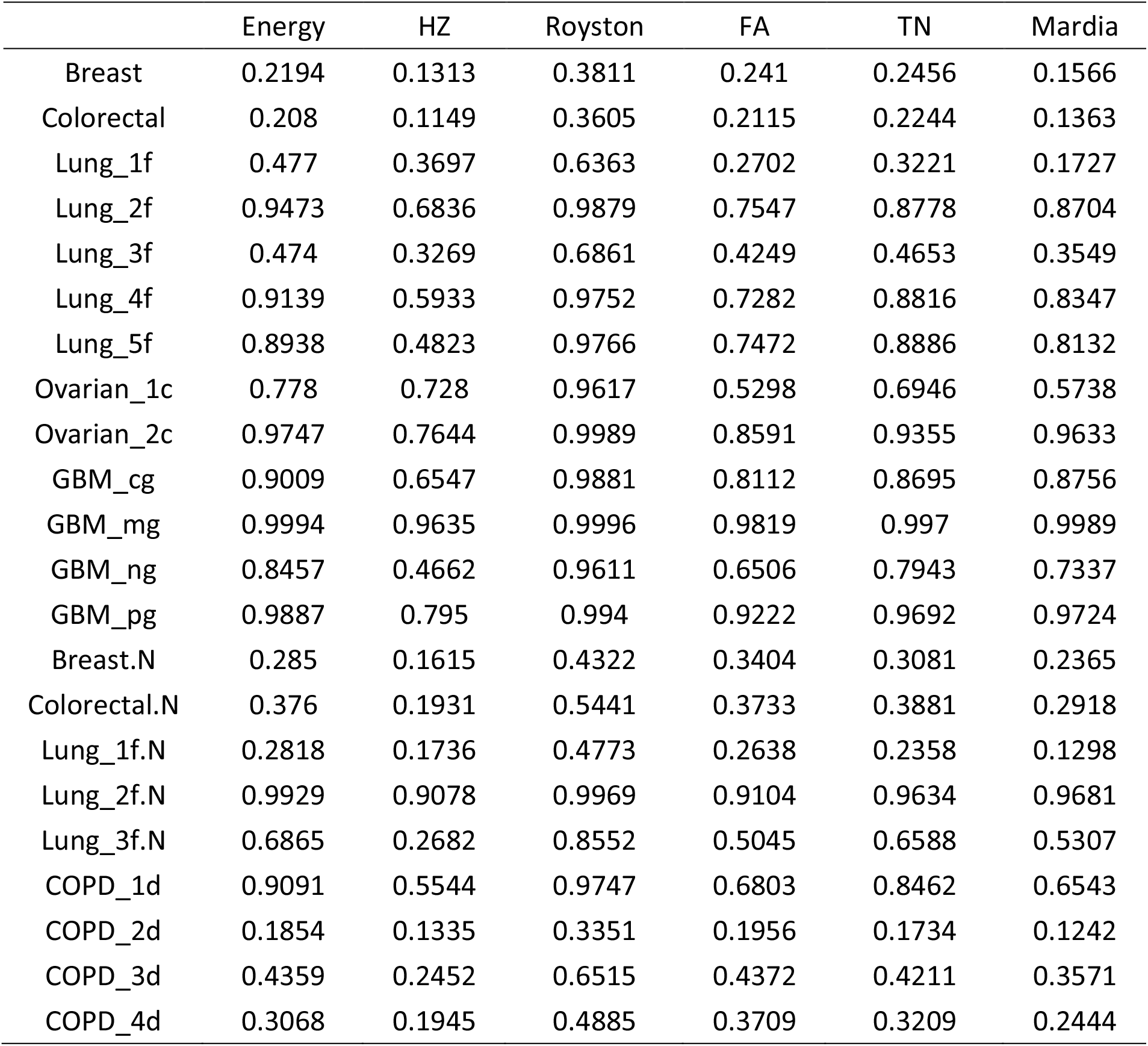
Average proportions *Q* across ten signaling pathways for each dataset with respect to each of the MVN test.

**Table S8:**
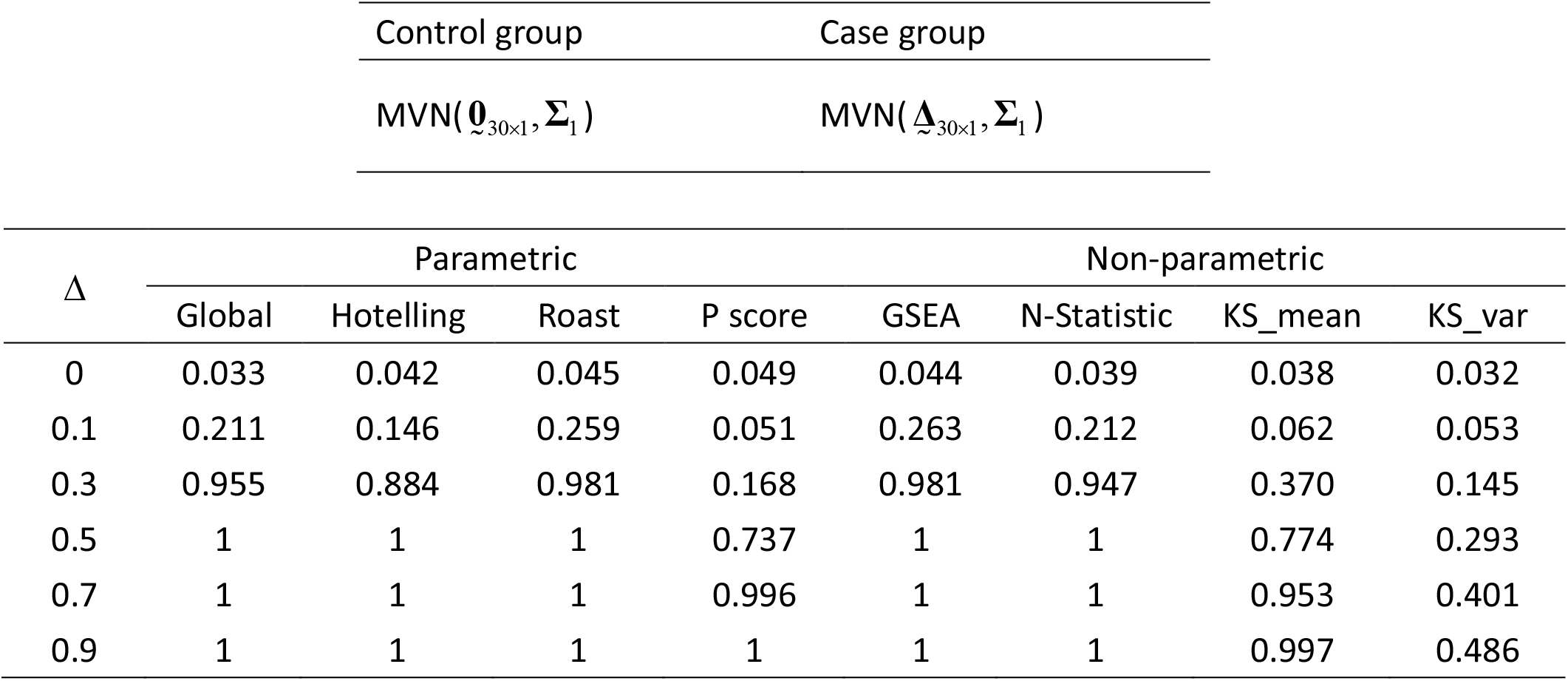
Proportion of rejection for each of the GSAs under the following distributions

**Table S9:**
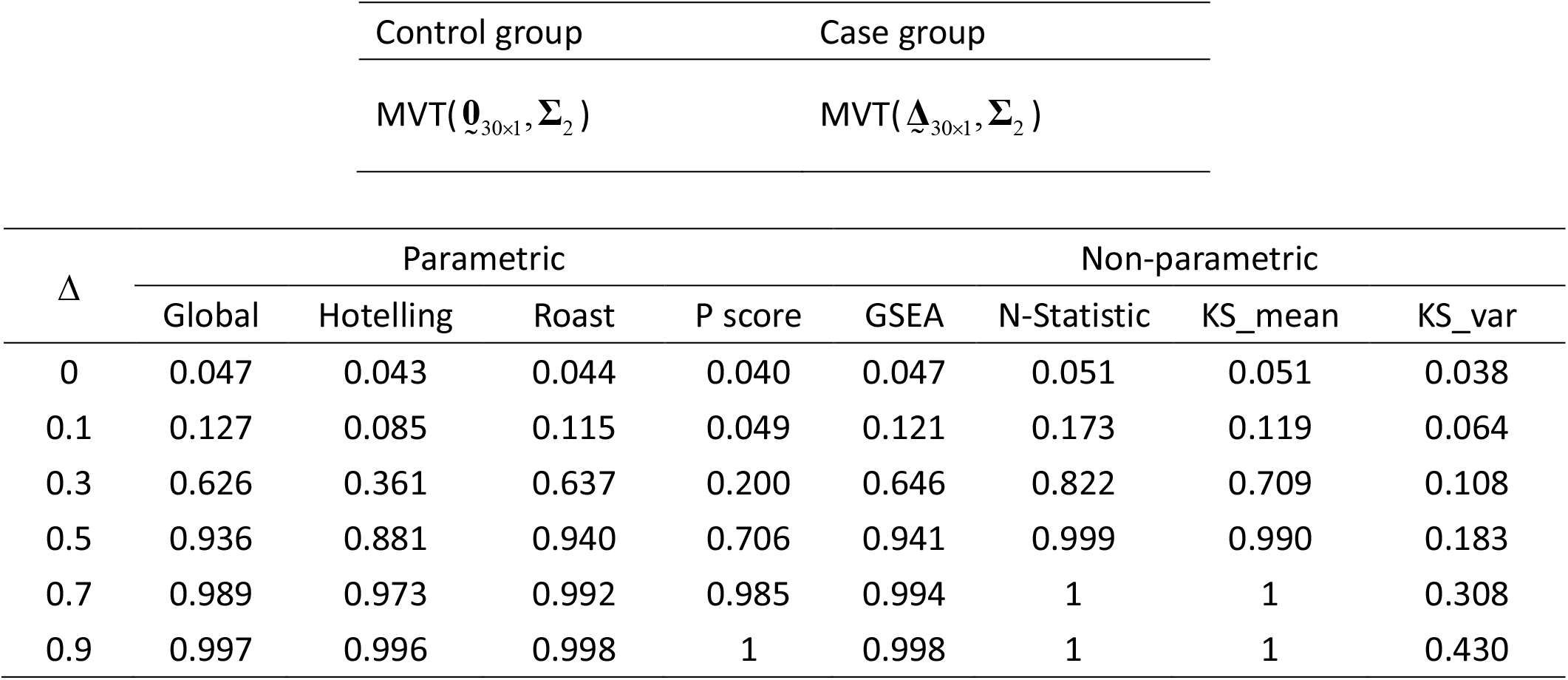
Proportion of rejection for each of the GSAs under the following distributions

**Table S10:**
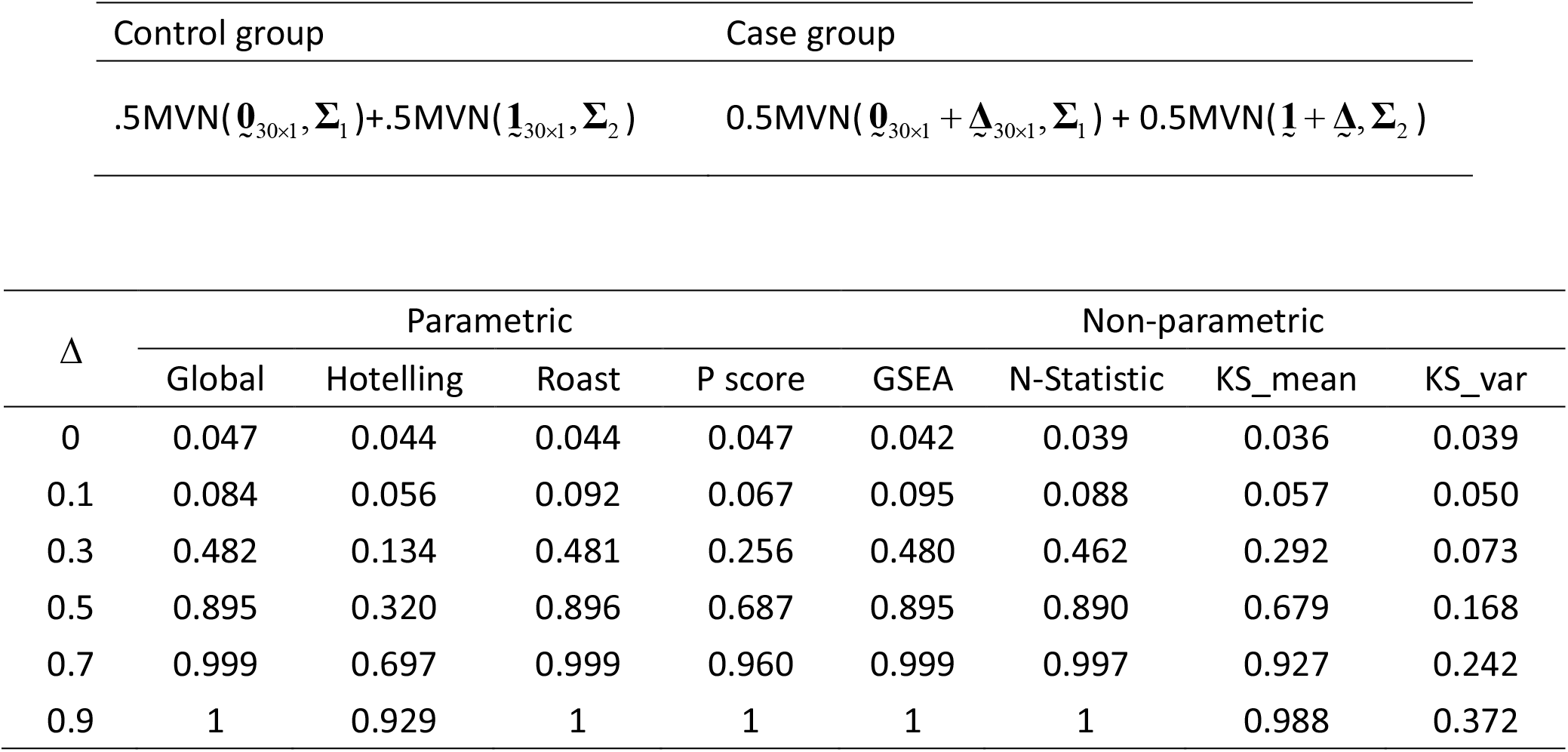
Proportion of rejection for each of the GSAs under the following distributions

**Table S11:**
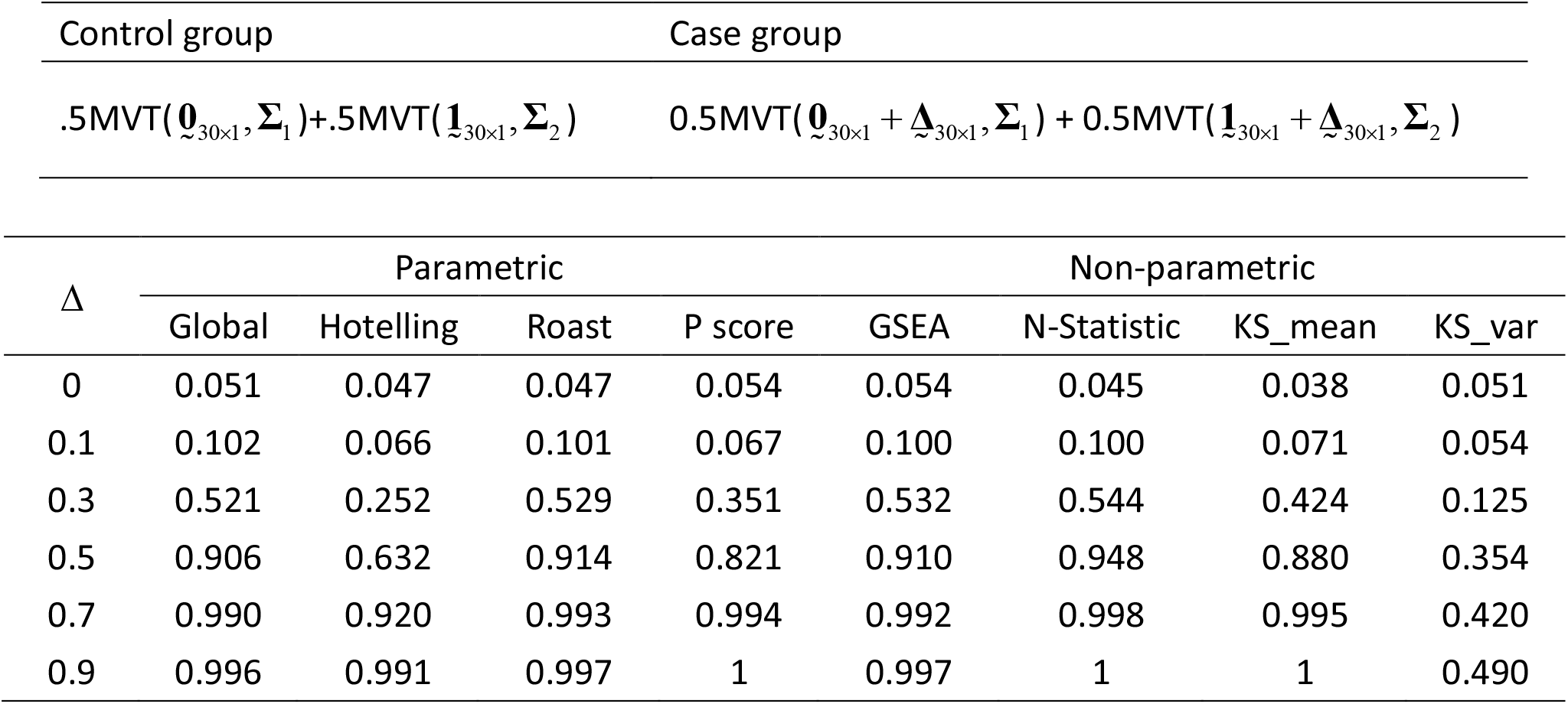
Proportion of rejection for each of the GSAs under the following distributions

**Table S12:**
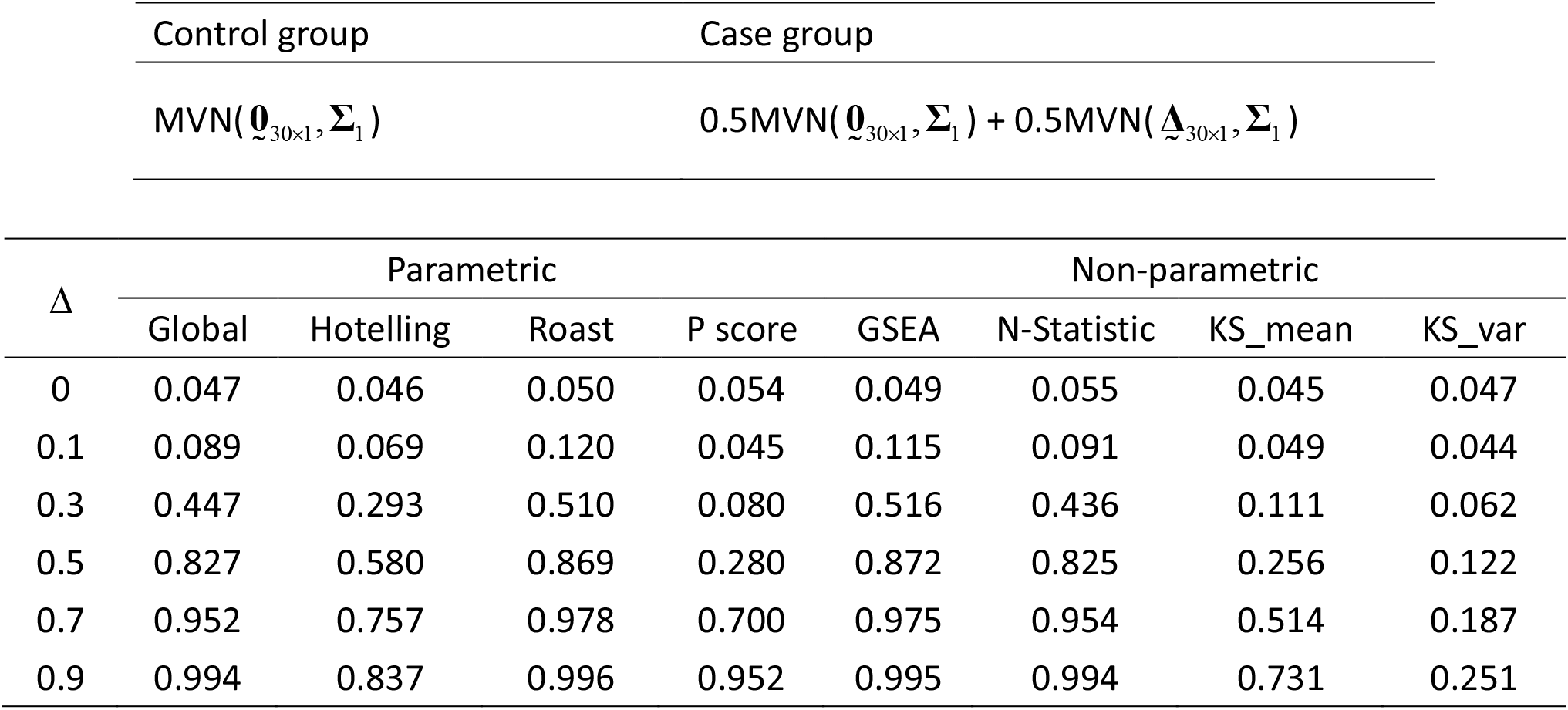
Proportion of rejection for each of the GSAs under the following distributions

**Table S13:**
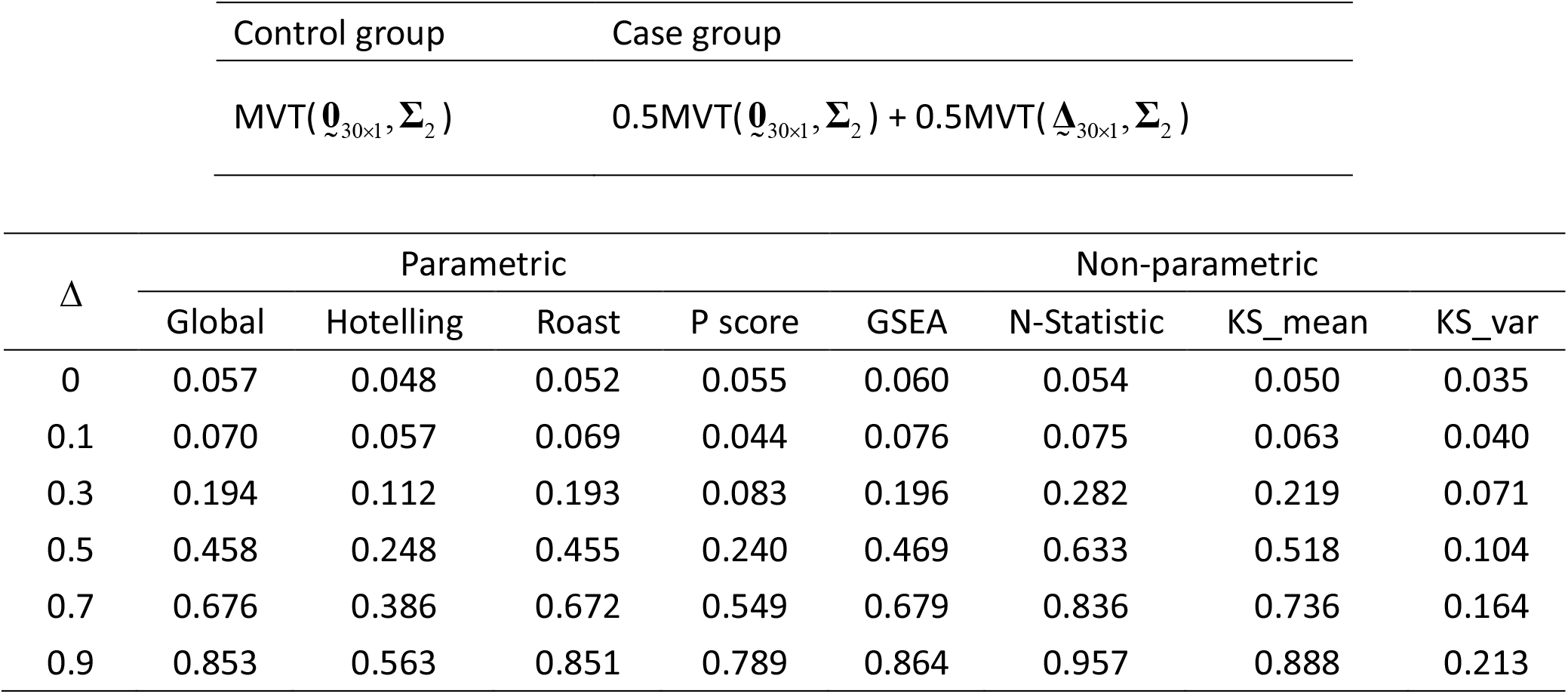
Proportion of rejection for each of the GSAs under the following distributions

**Table S14:** Proportion of rejection for each of the GSAs under the following distributions

| Control group |  | Case group |  |  |  |  |  |  |
| --- | --- | --- | --- | --- | --- | --- | --- | --- |
| $MVN(\mathbf{0}_{30 \times 1}, \mathbf{\Sigma}_1)$ | | $0.4MVN(\mathbf{0}_{30 \times 1}, \mathbf{\Sigma}_1) + 0.3MVN(0.5 \times \mathbf{\Delta}_{30 \times 1}, \mathbf{\Sigma}_1) + 0.3MVN(\mathbf{\Delta}_{30 \times 1}, \mathbf{\Sigma}_1)$ | | | | | | |
| $\Delta$ | Parametric | | | | Non-parametric | | | |
|  | Global | Hotelling | Roast | P score | GSEA | N-Statistic | KS_mean | KS_var |
| 0 | 0.043 | 0.036 | 0.044 | 0.054 | 0.043 | 0.040 | 0.040 | 0.046 |
| 0.1 | 0.084 | 0.060 | 0.096 | 0.047 | 0.104 | 0.081 | 0.066 | 0.035 |
| 0.3 | 0.345 | 0.207 | 0.417 | 0.071 | 0.421 | 0.339 | 0.078 | 0.053 |
| 0.5 | 0.701 | 0.469 | 0.793 | 0.166 | 0.792 | 0.691 | 0.198 | 0.087 |
| 0.7 | 0.938 | 0.732 | 0.967 | 0.493 | 0.967 | 0.941 | 0.392 | 0.161 |
| 0.9 | 0.986 | 0.853 | 0.992 | 0.840 | 0.992 | 0.985 | 0.599 | 0.228 |

**Table S15:** Proportion of rejection for each of the GSAs under the following distributions

| Control group |  | Case group |  |  |  |  |  |  |
| --- | --- | --- | --- | --- | --- | --- | --- | --- |
| $MVT(\mathbf{0}_{30 \times 1}, \mathbf{\Sigma}_2)$ | | $0.4MVT(\mathbf{0}_{30 \times 1}, \mathbf{\Sigma}_2) + 0.3 MVT(0.5 \times \mathbf{\Delta}_{30 \times 1}, \mathbf{\Sigma}_2) + 0.3 MVT(\mathbf{\Delta}_{30 \times 1}, \mathbf{\Sigma}_2)$ | | | | | | |
| $\Delta$ | Parametric | | | | Non-parametric | | | |
|  | Global | Hotelling | Roast | P score | GSEA | N-Statistic | KS_mean | KS_var |
| 0 | 0.048 | 0.042 | 0.047 | 0.054 | 0.041 | 0.043 | 0.037 | 0.053 |
| 0.1 | 0.076 | 0.053 | 0.074 | 0.047 | 0.079 | 0.074 | 0.070 | 0.047 |
| 0.3 | 0.185 | 0.109 | 0.191 | 0.070 | 0.196 | 0.248 | 0.199 | 0.068 |
| 0.5 | 0.388 | 0.190 | 0.377 | 0.150 | 0.390 | 0.505 | 0.405 | 0.070 |
| 0.7 | 0.602 | 0.339 | 0.610 | 0.400 | 0.622 | 0.781 | 0.676 | 0.144 |
| 0.9 | 0.777 | 0.445 | 0.774 | 0.648 | 0.789 | 0.923 | 0.825 | 0.151 |

